# Hidden Rice Diversity in the Guianas

**DOI:** 10.1101/562769

**Authors:** Tinde van Andel, Margret Veltman, Alice Bertin, Harro Maat, Thomas Polime, Derk Hille Ris Lambers, Jerry Tjoe Awie, Hugo de Boer, Vincent Manzanilla

## Abstract

Traditional crop varieties are an important source of genetic diversity for crop adaptation and modern breeding. Landraces of Asian (*Oryza sativa*) and African (*Oryza glaberrima*) rice have been well studied on the continents where they were domesticated. However, their history of cultivation in northern South America is poorly understood. Here we reveal the rice diversity that is maintained by Maroons, descendants of enslaved Africans who fled to the interior forests of the Guianas ca. 300 years ago. We interviewed subsistence farmers who practice shifting cultivation along the Maroni and Lawa rivers that form the natural border between French Guiana and Suriname, and used ethnobotanical and morphological methods to identify around 50 varieties, of which 15 were previously undocumented. The genetic origin of these varieties was explored using the Angiosperms-353 universal probe set. Despite the large distances between sites and relative inaccessibility of the area, phenotypic and genetic diversity did not display any geographic structure, which is consistent with knowledge of seed exchange among members of the same ethnolinguistic group. Although improved US cultivars were introduced in Maroon villages in the 1940s, these have not displaced the traditional landraces, which are cherished for their taste and nutritious qualities and for their importance in Maroon spiritual life. The unique agricultural and ritual practices of Maroons confirm their role as custodians of rice diversity, a role that is currently under threat from external pressures and encroaching globalization. We expect that the rice diversity uncovered in this study represents only a fraction of the total diversity in the Guianas and may constitute a large untapped resource that holds promise for future rice improvement. Further efforts to inventory and preserve these landraces will help to protect a precious cultural heritage and local food security.

## 1 Introduction

Rice is the most widely consumed staple food in the world. Two species of domesticated rice exist: the widely cultivated Asian rice (*Oryza sativa* L.), domesticated in China some 10,000 years ago, and the lesser-known African rice (*O. glaberrima* Steud), domesticated about 3,000 years ago (Stein et al., 2018). There are thousands of rice cultivars in Asia, Africa and the Americas, and a significant proportion of this diversity is maintained in germplasm institutes (Jackson and Lettington, 2002; Sanchez et al., 2013). With a growing world population and increasing impacts of climate change, rice breeders urgently need to develop more sustainable cultivars with higher yields, healthier grains and reduced environmental footprints (Stein et al., 2018; Wang et al. 2018). The narrow genetic base of modern crop cultivars guarantees phenotypic uniformity and genetic stability, but also makes them vulnerable to environmental fluctuations, such as climate change, reduced soil fertility, pests and diseases (Zeven, 1998).

Wild relatives of rice and traditional landraces often show adaptations towards marginal environments and pest resistance and are therefore considered as an untapped genetic resource for breeding new cultivars resilient to future challenges (Alvarez et al., 2007; Wang et al. 2018). Landraces are also of key importance in local food security and preserving cultural heritage (Perales et al., 2005; Ardenghi et al., 2018), and reveal past migration patterns of humans and their contacts with outsiders (Westengen et al., 2014). Over the last few decades, a severe genetic erosion of crops has taken place due to the replacement of landraces by modern cultivars. Much effort has been made to safeguard landraces *ex situ* (in gene banks) to make their genetic resources available for breeders, but the *in situ* conservation of agrodiversity within traditional farming systems has not been pursued to the same extent (Maxted et al., 2002). Ethnobotanical inventories are powerful tools in detecting these neglected genetic resources and understanding the social and cultural factors involved in generating and maintaining their diversity and distribution (Westengen et al., 2014; Ardenghi et al. 2018). Here we describe the rice landraces that are grown by Maroons, descendants of escaped slaves in Suriname and French Guiana and discuss their efforts and motivations to maintain this diversity.

Rice has been grown for centuries in the Guianas (Guyana, Suriname and French Guiana). In the 17th and 18th century, plantation owners imported the crop from the US and West Africa as provision for their enslaved laborers (Carney, 2009; Van Andel et al. 2016a). Asian rice was introduced by Portuguese sailors in West Africa in the 16^th^ century, and both African and Asian rice were grown by African farmers before the onset of the transatlantic slave trade (Linares, 2002). Rice was sold in the husk, allowing for longer storage and germination ability (Carney, 2009). After their arrival in the Americas, slaves managed to gather leftover seeds from ship’s hold or other storage places and planted these in small provision grounds at the periphery of the plantations (Price, 1991; Van Andel et al., 2012). These home gardens enabled them to grow at least some of the familiar crops of their motherland and became central to their physical and spiritual life. The rice they cultivated consisted most likely of upland, rainfed varieties (Carney, 2009; Fleury, 2012). When they escaped from the plantations, Maroons brought crop seeds and established new gardens in their communities along the major rivers in the forested interior of Guyana, Suriname and the French side of the Maroni River (Codd and Peterkin, 1933; Hurault, 1965; Price, 1991). Today, there are no Maroons left in Guyana, but six ethnic groups of Maroons still remain in Suriname and French Guiana: Paramaccans, Aluku (or Boni), Kwinti, Matawai, Ndyuka (or Aucans) and Saramaccans (Price, 2013). Rice has been their staple food for centuries, enabling them to survive in the remote hinterland, independent from coastal societies (Price, 1993; Fleury, 2012). In 2013, Maroons numbered some 210,000 people and constituted around 25% of the population of Suriname and French Guiana (Price, 2013).

Commercial rice cultivation in the Guianas started only in the early 1900s in the coastal swamps of Guyana and Suriname. The first improved cultivars were based on traditional varieties brought by indentured laborers from India and Java, who were recruited to work on the plantations after slavery was abolished (Codd and Peterkin, 1933; Stahel, 1933). From the 1950s onwards, commercial Asian wetland cultivars suitable for mechanical harvesting were planted in newly constructed polders in Suriname (Young and Angier, 2010; Maat and van Andel, 2018). Commercial rice fields in French Guiana were not developed until 1982, with the construction of coastal polders around Mana (Clément et al., 2011). Today, new cultivars continue to be developed by the Anne van Dijk Rice Research Centre (SNRI/ADRON, www.snri-adron.com), the national institute for rice research in Nieuw Nickerie, Suriname.

Traditional Maroon agriculture is under pressure by increasing commercial and governmental interventions. Gold mining and logging concessions have been issued on traditional territories, while prospects of education and employment stimulate migration to urban areas (Paramaribo, Cayenne, St. Laurent), which in turn leads to a shortage of farm labor (Heemskerk, 2000; Fleskens and Jorritsma, 2010, Price, 2012). Recent infrastructural developments (roads, airstrips) have facilitated access to the remote hinterland (Price, 2012) and commercially produced rice is now widely available in gold miner shops in the interior (Heemskerk, 2000). Shortening of fallow periods in shifting cultivation plots and limited use of fertilizers keep soil productivity low, with yields rarely exceeding 1000 kg/ha and arguably not meeting local demand (Baumgart et al., 1998; Fleskens and Jorritsma, 2010; Nascente and Kromocardi, 2017). The increasing influence of evangelical churches in Maroon territory, development organizations and government policies have resulted in the perception of traditional Maroon farmers as ‘backward’ and in need of modernization (Fleskens and Jorritsma, 2010; Heemskerk, 2003; Léobal, 2016). These factors actively discourage traditional Maroon farming, and put their landraces at risk of disappearing.

No systematic inventory of Maroon rice landraces with storage of vouchers or germplasm has been made thus far and their varieties have not been included in formal breeding experiments or field trials. In 1936, a French agronomist discovered more than 30 rice varieties cultivated by Maroons along the Maroni and Tapanahoni rivers (Vaillant, 1948). Almost 20 years later, Portères (1955) identified these collections as *Oryza sativa*, except for one sample that represented African rice (*O. glaberrima*). Vaillant (1948: 520) made an urgent appeal to continue the study of Maroon rice varieties, as they might ‘represent a detached branch of historical African cultivars at a time where European culture and importation of selected hybrids had not yet played a role’, but this was not followed up. Anthropologists Hurault (1965), Bilby et al. (1989) and Price (1993) mentioned Maroon rice diversity, but hardly published any landrace names. Biologist Geijskes (1955) recorded 23 names of rice varieties along the Maroni River, and anthropologist Fleury (2016) listed 21 names among the Aluku on the Lawa River, but no herbarium or seed collections were made. The only expedition to collect Maroon rice varieties was carried out by Baumgart et al. (1998) along the upper Suriname River, who deposited their samples at the SNRI/ADRON seed bank. The discovery of a field of *O. glaberrima* in a Saramaccan village in 2008 (Van Andel, 2010), stimulated further ethnobotanical research on African crops grown among Maroons (Van Andel et al. 2016b) and rice in particular (Van Andel et al. 2016a; Maat and Van Andel, 2018), but genetic analysis of rice was limited to one accession.

To complement these previous anthropological and ethnobotanical studies, plants can now be studied through the lens of genomics. Next generation sequencing analysis can reveal further details about the origins, evolution and diversity of different crop varieties. Specifically, universal probes have recently been designed for target enrichment of 353 low copy nuclear genes across the Plant and Fungal Trees of Life (PAFTOL) (Johnson et al., 2018). Target enrichment is a versatile, reproducible and scalable technique that increases the representation of selected portions of the genome prior to its sequencing (Lemmon and Lemmon, 2013). The Angiosperms-353 probe set allows the generation of high-quality genomic data for all the angiosperm taxa, making results comparable at an unprecedented taxonomic breadth, but its utility for differentiating closely related samples of the same species has not yet been demonstrated.

In this study, our first aim was to collect herbarium vouchers, seed material and DNA samples of different rice species and landraces grown by Maroons along the French Guiana-Suriname border. Our second aim was to document their morphological characters and associated traditional knowledge on their agricultural requirements, cultivation and processing methods, local names and meanings, culinary and spiritual values. Based on previous studies, we anticipated that Maroon rice varieties would be known under a range of local names and hard to tell apart visually. We therefore tested the ability of the Angiosperm-353 markers to capture the genetic diversity of these traditional landraces and compared this with other commercial cultivars and traditional varieties grown in the region, demonstrating the functionality of the Angiosperms-353 universal probe set (Johnson et al., 2018) for population genomics.

## 2 Materials and Methods

### 2.1 Ethnobotanical survey

Before the start of the fieldwork, a prior informed consent form was prepared in French following the guidelines of the Collectivité Territoriale de Guyane in Cayenne. This document was signed on 10 April 2017 by Mr. Jacques Chapel Martin, captain of the village of Grand Santi and responsible for the Ndyuka traditional authorities of the region Grand Santi (Lawa River). St. Laurent du Maroni is considered an urban area where no traditional authorities are in place (Tareau et al., 2017), so no such document was needed. Fieldwork was done from 4 to 22 July 2017. We interviewed Maroon rice farmers around St. Laurent and Bigiston (lower Maroni River), near Providence (upper Maroni River), near the Gonini River mouth, and around Grand Santi (Lawa River), located ca. 144 km upstream from St. Laurent (Figure 1). These locations were chosen because of their relatively easy access and previous contacts with local Maroon farmers.

**Figure 1.**
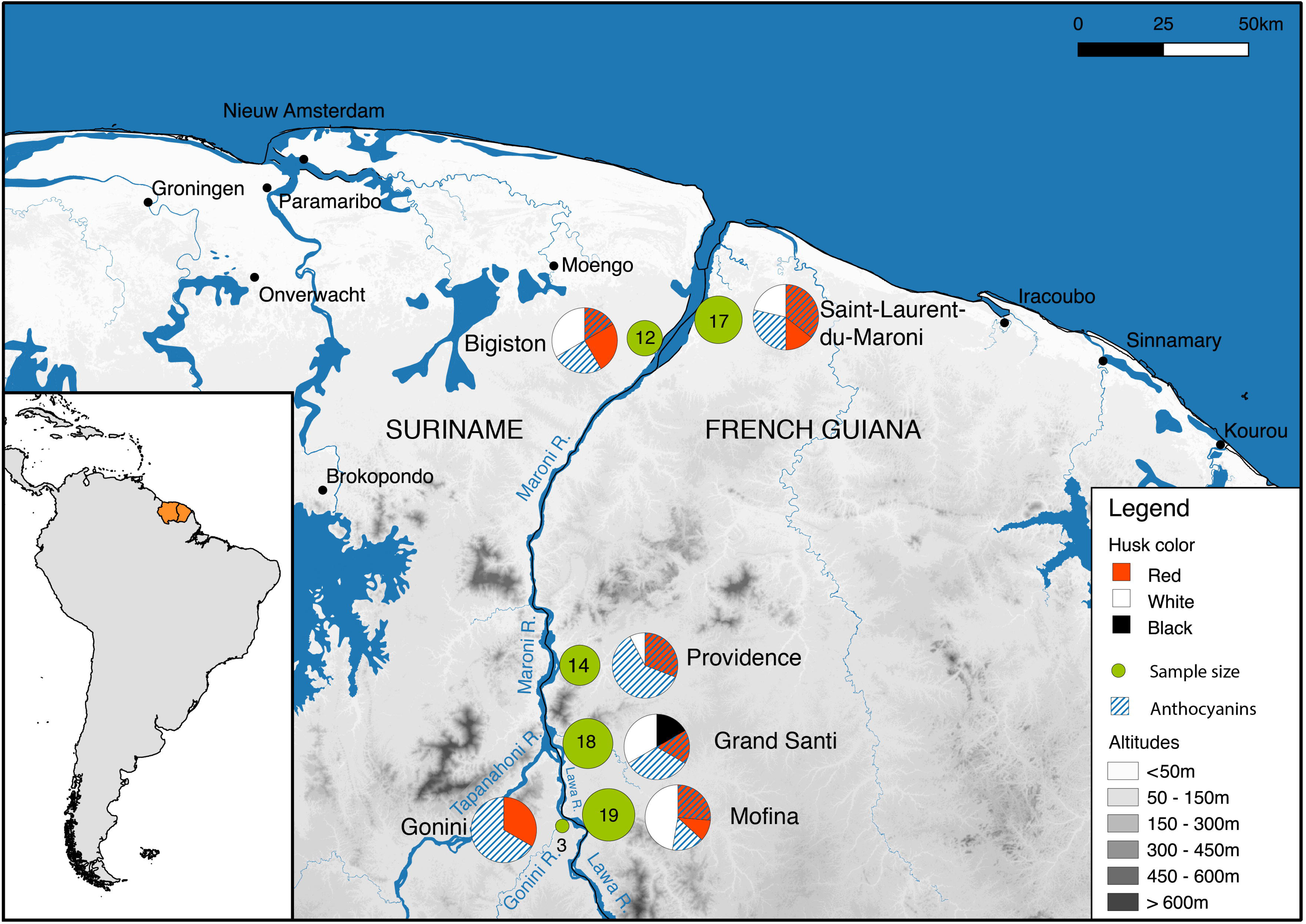
Study area. Fieldwork locations are represented with green circles, whose size is proportional to the number of samples. The pie charts illustrate the proportion of samples with different husk colors and anthocyanins. Metadata for this map are provided in Supplementary Table S3.

Local rice farmers were recruited by our translators, who explained our research objectives in the Ndyuka or Saramaccan language if they were not fluent in Dutch, French or Sranantongo (the *lingua franca* in Suriname). Having obtained the farmers’ oral prior informed consent, we conducted face-to-face interviews and visited their rice fields if they had a standing crop. Herbarium vouchers of living rice plants were collected using standard botanical methods, and one duplicate was deposited at the Herbier IRD du Guyane (CAY) in Cayenne, French Guyana and the other in the herbarium of Naturalis Biodiversity Center (L) in Leiden, the Netherlands. When no living plants were available for specific varieties, we collected seeds from rice stored in people’s outdoor kitchens. Seed samples were stored in paper envelopes: one duplicate of each variety was sent to the SNRI/ADRON germplasm bank in Suriname for storage and phenotyping, while the other was stored at the Economic Botany collection of L. We used the ‘rice passport data table’, developed by the SNRI/ADRON, to document local names, agronomical and culinary characteristics of each collected rice variety. We expanded this table into a semi-structured interview by adding questions concerning positive and negative features of landraces (Supplementary Table S1). We also posed additional questions about challenges faced in rice cultivation, the use of agrochemicals and the importance of rice in ancestor rituals.

For each collected rice variety, we documented morphological characteristics in the field (size, husk and pericarp color, presence of awns, anthocyanins, etc.) and made detailed pictures of living plants, panicles, husked and dehusked seeds. To facilitate the discussion in the field, we made a ‘rice book’: a ring binder with samples of local rice varieties secured under transparent tape (Supplementary Figure S1). We made short videos on the different stages of rice cultivation and processing and uploaded these on youtube (Supplementary Material). To raise awareness on local rice diversity, we designed a poster with pictures of the different rice varieties and associated traditional information, which will be distributed in Suriname and French Guiana (Supplementary Figure S2). Throughout this paper, we follow Zeven (1998) and use the term ‘landrace’ for farmer-developed rice accessions, ‘cultivar’ for rice accessions developed by companies or breeding institutes and ‘varieties’ for all rice accessions, regardless of genetic improvement.

### 2.2 Genetic analysis

Fresh leaf material from living rice plants were stored on silica gel. In five cases, we germinated seeds on wet tissue paper and extracted DNA from the green sprouts. DNA was extracted from approximately 40 mg of dry leaf material using the DNeasy Plant Mini Kit (Qiagen). Total DNA (0.2-1.0 μg) was sheared to 500 bp fragments using a Covaris S220 sonicator (Woburn, MA, USA). Dual indexed libraries were prepared using the Meyer and Kircher protocol (Meyer and Kircher, 2010) for shotgun sequencing and target capture. We captured target sequences with the PAFTOL project probe set (Johnson et al., 2018). We prepared and pooled 41 equimolar libraries in two capture reactions with an average 300 ng of input DNA per pool. The RNA probes were hybridized for 24 hours before target baiting, and 14 PCR cycles were carried out after enrichment following the MyBaits v.3 manual. The enriched libraries libraries were sequenced on one Illumina HiSeq 3000 lane (150bp paired-end).

To find out whether the Maroon rice varieties bear genetic similarity to other Asian rice grown in our study area, we included ten accessions from French Guiana and Suriname for which genetic data is freely available (Supplementary Table S2). These accessions were resequenced as part of the 3000 Rice Genomes Project (3K RGP) and represent a mix of populations and subspecies, including *Oryza sativa* ssp*. japonica* (4), *O. sativa* ssp. *indica* (4) and two admixed individuals between *O. sativa* ssp*. japonica* and *O. sativa* ssp. *indica* (Wang et al., 2018). The raw sequencing reads were trimmed and quality filtered using Trimmomatic v.0.32 (Bolger et al., 2014) and mapped with the Burrows-Wheeler Aligner v.0.7.5a BWA-MEM algorithm (Li and Durbin, 2009) against the reference genome of *Oryza sativa* subsp*. indica* (ref. ASM465v1). The BAM files were filtered to a minimum quality score of 30 and read depth of 5 with samtools v.1.3.1 (Li et al., 2009). SNPs were called with ANGSD v.0.549 (Korneliussen et al., 2014) and filtered with vcftools v.0.1.13 (Danecek et al., 2011). Sites with more than 20% missing data were discarded. SNPs were pruned with PLINK v.1.90b5.2 (Purcell et al., 2007) using a pairwise correlation threshold of 0.5 in sliding windows of 500 SNPs with step size 50.

Multi-dimensional scaling of the genetic variation was performed in PLINK v.1.90b5.2. Population structure analyses were conducted with ADMIXTURE v.1.3 (Alexander and Lange, 2011). Genetic differentiation between sampling locations was estimated with Weir and Cockerham’s weighted Fst statistics (Weir & Cockerham, 1984) as implemented in vcftools v.0.1.13. The average genetic diversity per population was calculated as π in vcftools and tested for significance with the non-parametric Kruskal-Wallis test. To ensure independent measures, we chose a window size of 150 kb, which exceeds the average extent of linkage disequilibrium as estimated in *O. sativa* ssp. *japonica* (Mather et al., 2007). An approximately maximum-likelihood (ML) tree was constructed in FastTree v.2.1 (Price et al., 2010), after removing heterozygous sites with bcftools v.1.1 (Li et al., 2009) and concatenating SNPs with a custom perls script (Bergey, 2012). Pairwise genetic distances were computed using the p-distance model in MEGA7 (Kumar et al., 2016). All scripts and command line options used for the analyses are openly available on Zenodo (10.5281/zenodo.2560038).

## 3 Results

### 3.1 Farming practices

We interviewed 21 rice farmers, 20 of Ndyuka and one of Saramaccan descent, of which eight in the lower Maroni region (St. Laurent, Bigiston, Manjabon), three near Providence (upper Maroni River), six in Grand Santi and four near the Gonini River mouth (Figure 1). All participants were female except one, who showed us the field of his sister-in-law. No male rice farmers were mentioned, as it is the Maroon women who do the planting, weeding, harvesting and processing of rice, either by themselves or together with female relatives. Men do help with the clearing and burning of fields in the dry period around November. Rice is sown as the first crop in the fertile ashes at the onset of the short rainy season between December and January. After weeds and debris have been removed by hand, rice is directly sown in the soil, either in small holes (‘diki olo’) dug with a hoe (‘tyap’), or cast (‘fringi’) over a tilled field (Supplementary Material, video 1). Rice grains must be covered by loose soil and pressed flat to protect them from birds that feed on exposed seeds. One farmer assured us that birds in primary forest are not yet accustomed to rice fields, making it is possible to sow the grains without covering them. Most rice is sown in well-drained, sandy upland soils on fields cut in primary forest or in secondary forest in various stages of succession. Three of the interviewed farmers cultivated rice on brown clay soils in a floodplain forest, which was very successful, while one farmer grew her crop in a swampy white sand savanna, with meager results. In Grand Santi (French Guiana), some women said they do not go to their rice fields on Fridays, as this is prohibited by the ‘local God of the forest’. Therefore, some people also have a rice field on the Surinamese side of the Lawa River, as there the bush spirits forbid entry on Sundays. This way, they can work on their rice fields every day of the week. Given the numerous islands, unclear borders and absence of border control, many Maroons (temporarily) live and practice agriculture on both sides of the Lawa and upper Maroni rivers.

Generally, rice is harvested between March and May. In St. Laurent, other fields are then directly cut and burned, so a second rice crop can be harvested from July to early September. After this, no more rice is planted until December. In older fields, the rice grains that shatter during the first harvest from March to May will sprout between the cassava plants and mature four months later. Because of the lack of nutrients, this ‘grow-back rice’ has meager-looking stems with few-seeded panicles. The crop is also much shorter (80-90 cm) than rice sown on freshly burned fields (>1.60 m), but harvested nevertheless. Farmers in Grand Santi do not plant a second rice field after their first harvest in April, as there is too much rain in July for the rice to mature. Precipitation data (World Bank Group, 2018) indeed show a slightly higher rainfall in July in Grand Santi (198 mm) than in St. Laurent (186 mm). Since no mature rice fields were present around Grand Santi during our fieldwork in July, we made our herbarium collections in this region from ‘grow-back rice’.

Farmers sow their varieties one after another on different parts of their fields and harvest them sequentially to divide the workload over time and facilitate separate storage. None of the farmers we interviewed used any pesticides or fertilizers. While most varieties mature within four months, the white Alekisola, Watralanti and Mesti take longer, so they are often sown first and harvested last. The ripe ears are cut off with a small knife (Figure 2, Supplementary Material, video 2), tied to sheaves (Figure 3), and dried for some time in the sun on corrugated iron sheets or woven polypropylene rice bags. When fully dry, the bundles are stored separately per variety in plastic buckets, steel oil drums, glass bottles or rice bags. According to the farmers, some varieties take longer to dry than others. Freshly picked rice is sometimes consumed the same day, but then the ears are dipped in boiling water to facilitate removal of the husks. Farmers said that when still in their tough, fibrous husk, rice seeds can be stored for as many as three years without spoiling or losing their ability to germinate. Sowing stock (‘paansu’) is not stored separately from the rice meant for consumption.

**Figure 2.**
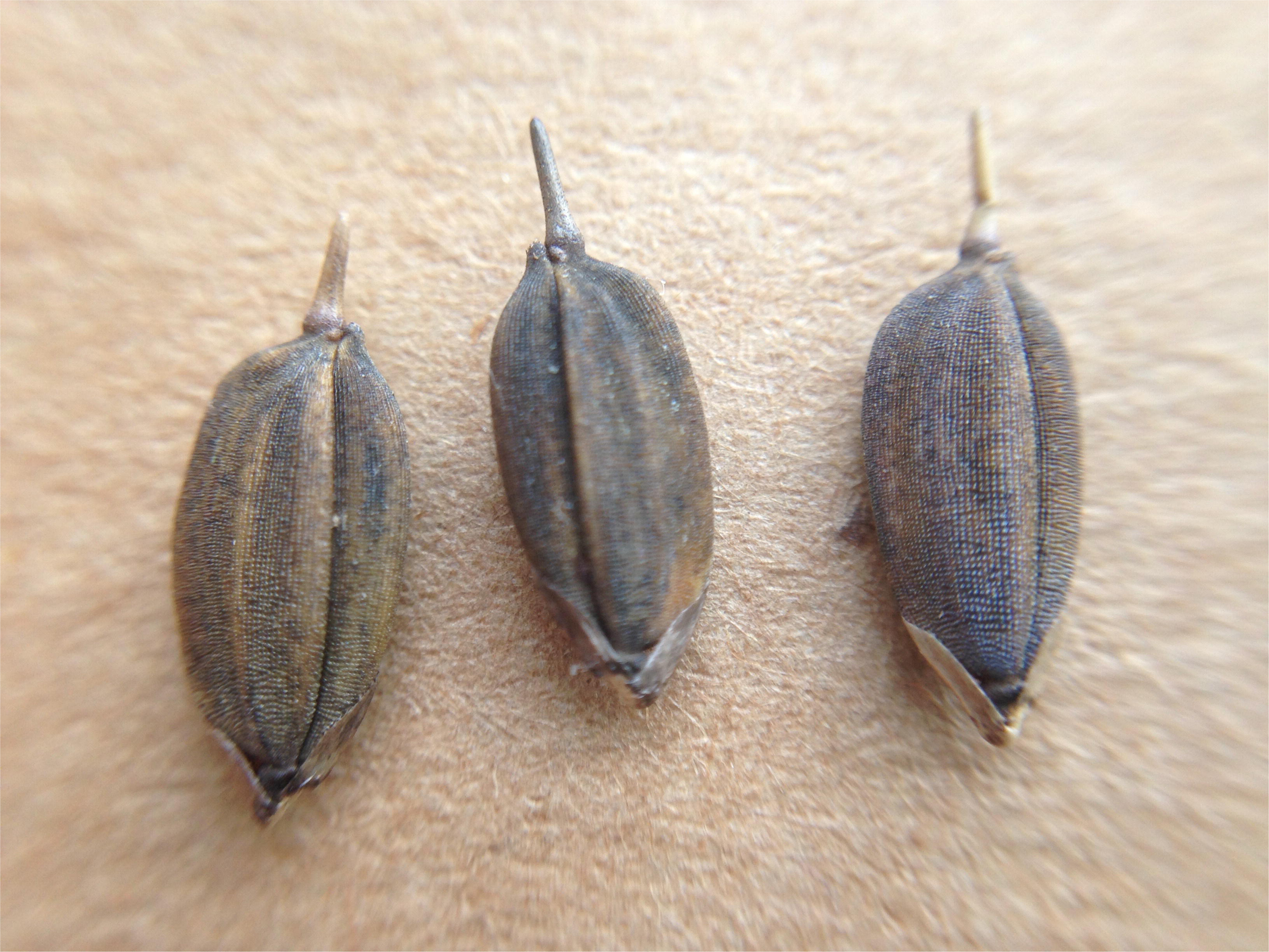
Harvesting ripe panicles by hand near Bigiston, lower Maroni River. Picture by Alice Bertin.

**Figure 3.**
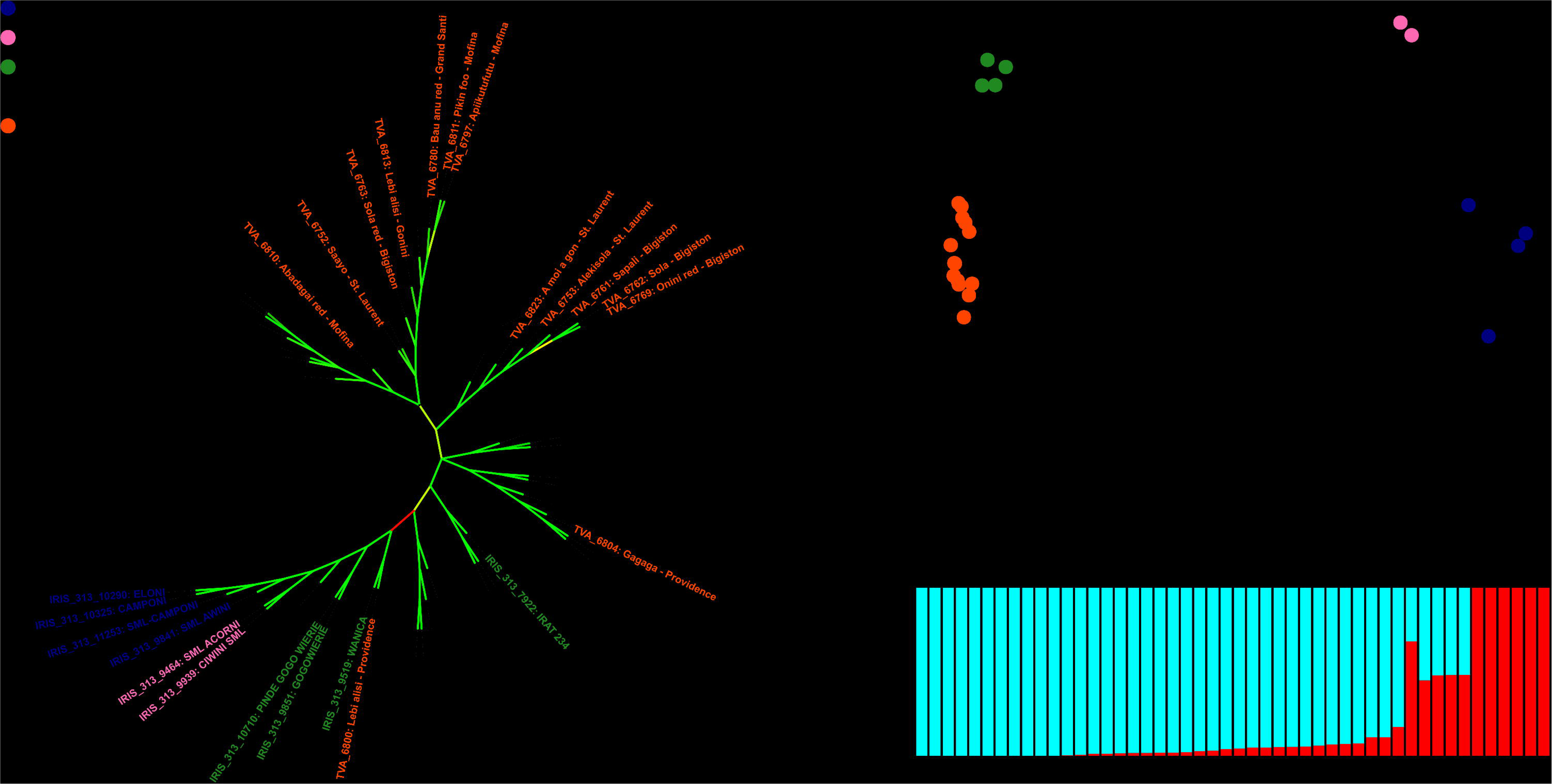
Dried rice tied into sheaves, St. Laurent. In the background a plastic drum in which dried rice is stored for next year’s planting stock and consumption. Picture by Alice Bertin.

Threshing is done before storage (when seeds are stored separately in oil drums) or when it is time to consume the rice. Bundles are spread out on a plastic sheet and the seeds are separated from the ears by spreading out the sheaves and trampling on them barefoot with a dancing movement (Supplementary Material, video 3). Rice varieties with scabrous stems are sometimes threshed by putting the bundles in a bag and beating it with a stick. Only just before consumption, the rice is hand dehusked by pounding it with a pestle in a heavy wooden mortar Supplementary Material, video 4). To winnow the rice, it is put on a large wooden tray and thrown into the air to let the wind blow away the empty husks (Supplementary Material, video 5). The pounding process can be repeated to also remove the bran (pericarp), and get ‘clean and very white rice’, with broken grains, if this is preferred. Some villages have diesel-powered rice mills, for which the farmers have to pay € 4 to mill a 20-liter tin of rice.

### 3.2 Motivations for growing rice

A major motivation for growing rice is that it is much cheaper than buying it in a store. In most Maroon families, rice is eaten with every meal. The farmers we interviewed said that with a good harvest, they would have sufficient rice to last for an entire year or to the next harvest. Some women kept more than two full oil drums (200 liter each) of rice grains in stock in their outdoor kitchen. A few occasionally sell rice to other Maroons who do not grow (enough) rice to be self-sufficient. On the ‘Ndyuka market’ on Charbonnière Plage in St. Laurent, 2-kg bags of milled Maroon rice were sold for € 5 per bag. Most farmers said they only buy rice when their own stocks are depleted. Since most shopkeepers in the interior of Suriname and French Guiana are Chinese entrepreneurs, shop rice is known as ‘Chinese rice’, although it is commercially produced wetland rice from Nickerie (mostly ‘Breuk III’, a cheap type of broken rice of the Manglie brand). A 25-kg bag of this broken rice sells in the interior for € 20, which is more than twice the price paid in Paramaribo (€ 7-8).

Almost all farmers we interviewed said that their home-grown rice is healthier and tastier than the type that is sold by local shops. People complained that ‘Chinese rice’ tasted ‘horrible’, often had sand or stones in it, could not be stored long, and was frequently infested by beetles or larvae. They also warned us that ‘eating only shop rice will give you disease’. Farmers repeatedly assured us that Maroon rice ‘has vitamins and makes your belly feel full’, has much ‘goma’ (starch) and ‘gives good milk’, referring to the foam produced by the bran of home-grown rice during cooking. One farmer remarked that the increasing number of diabetes patients in St. Laurent should eat traditional rice instead of shop rice to control their disease. Milling traditionally-grown rice varieties by hand with a mortar and pestle removes the husk but not the bran and the germ, so it is in fact ‘whole-grain rice’, which is higher in fibres, lipids, proteins, minerals and vitamins than polished rice and gives a fuller feeling after consumption (Heinemann et al., 2005; Ryan, 2011). Farmers remarked that when they mechanically process their rice, it loses some of its taste, as the machine mill removes all the bran, ‘and that is the healthy part of the rice’. They still prefer mechanically milled home-grown rice over shop rice.

Locally grown rice has symbolic value too. Funerals are important social events among the Maroons and may last several days, so relatives from faraway are able to attend. When someone dies in a traditional Maroon village, the family members are required to bring home-grown rice to the mourning ceremony (Price, 1993). A gift of shop-bought rice or machine-milled traditional rice is not accepted. Before the burial takes place, women jointly pound and winnow the rice and prepare huge quantities of different rice dishes, a part of which is placed on a banana leaf as a sacrifice for the deceased, while the remainder is consumed (Fleury, 2012; Reijers, 2014). Attendance of funerals was frequently mentioned as a motivation for growing rice, even by those who identified themselves as ‘church people’.

All but three of the farmers we interviewed obtained their planting stock as a gift from family members. From the other three, one bought it in a neighboring village and two purchased it on the Charbonnière market. Most Maroons in St. Laurent and Grand Santi are migrants who obtained their rice varieties from relatives in traditional villages deep in the interior of Suriname, as far as Tyontyon island and Mpsuusu (Ndyuka territory, Tapanahoni River) and Masia Creek (Saramaccan territory, upper Suriname River). One variety (Aluku paansu) was exchanged with Aluku Maroons on the upper Lawa River. None of the farmers reported having bought seeds in Paramaribo or Cayenne.

### 3.3 Challenges in growing rice

Many of the rice farmers we interviewed were young women in their 20s or 30s, who said they enjoyed farming and were dedicated to continue the practices that their ancestors had passed on to them. They sometimes criticized others who ‘became too lazy’ to practice traditional agriculture or mill rice by hand because of the French child support payments that allowed them to buy rice in shops. None of the farmers complained about shortages of farm labor or a lack of suitable areas to burn a field. Birds are considered a minor nuisance and insect pests were not reported. Although most farmers have a rice field within walking distance from their home, some have fields that are located further away and can only be reached by boat. In spite of the high fuel costs, this is still considered economically feasible. The only farmer who complained about decreasing soil fertility had a field that was invaded by saplings of *Acacia mangium* Willd. This Australian tree was introduced to French Guiana in the 1970s to restore degraded mining areas but became invasive. As fire activates its germination, it quickly colonizes new agricultural fields (Delnatte and Meyer, 2012). Locally known as ‘Mira udu’ (ant wood), it dominates secondary forests in the outskirts of St. Laurent. Farmers said the only way to get rid of it is by ring-barking. We did not observe *A. mangium* elsewhere along the Maroni or Lawa River. A major challenge mentioned by Maroon farmers around St. Laurent is harassment by the French police. They said that during the dry season, the authorities fly small airplanes over the area to detect burning fields. Fines can be up to € 10,000, so some rice fields are made far from home to avoid discovery.

### 3.4 Diversity of rice varieties

We collected 38 herbarium vouchers of rice plants from 21 agricultural fields and 47 seed samples from living plants, stored and dried panicles or loose seeds. All collections are listed with their local name, collection number, locality and associated information in Supplementary Table S4. From the seed collections and herbarium vouchers with mature seeds, 56 seed samples were sent to SNRI/ADRON. Eventually, 48 were successfully germinated, grown into fertile adult plants and phenotyped by the SNRI/ADRON staff in July 2018. Based on morphological characters, genetic analysis and local names, we grouped our collections into ca. 50 varieties (Supplementary Table S4). The number of different rice varieties encountered per farmer (on the field and/or in storage) ranged between 1 and 7, with a mean of 3.2. Several farmers indicated that they had grown more varieties that year (up to 9), but some stock had been depleted. Most of the varieties were encountered only once. We extracted DNA from 40 rice individuals of different local varieties (Supplementary Table S4). Target sequences were captured from 35 herbarium vouchers and five germinated seed samples. One individual was removed from the analysis due to insufficient data. Following quality filtering, the median sequencing depth was 5.5X, with a range of 1.1-9.8X, resulting in a data set consisting of 39 individuals with 250,218 single nucleotide polymorphisms (SNPs).

#### 3.4.1 Species and subspecies identification

All but one sample belong to the Asian rice species *Oryza sativa*.The only exception is Baaka alisi (nrs. 6782 and 6773A), which represents a variety of African rice (*O. glaberrima*). Although people said African rice was not often cultivated, we encountered three farmers who kept small bags or glass bottles in stock with this conspicuously straight-awned rice with black husks and red pericarp (Figure 4). The use of African rice was shrouded in some mystery. Two farmers said that ‘the ancestors came with this rice, they brought it and ate it’. One farmer said that if you pound the black rice well enough, it becomes white (as it loses its red bran), so she would cook and eat it mixed it with her homegrown Asian rice. Other people ensured us that black rice is never eaten nor brought as a gift or prepared during funerals. The rice is mostly sold for a good price (± € 1 for a small bag of 40 gram on the Paramaribo market) to be used in traditional medicine. People cook the milled rice into porridge and drink the starchy water when they suffer from diarrhea. The husked seeds are also used in herbal baths to get into contact with ‘deep ancestor spirits’ during Afro-Surinamese rituals. Two of the farmers who grew black rice said they did not perform these rituals themselves, because they were ‘church people’. Baumgart et al. (1998) reported that black rice is only planted to keep the birds away from the rest of the crop, but this seems unlikely, as it is sold for a higher price than any of the other varieties.

The other varieties we collected were all identified as Asian rice and, according to the farmers, mostly upland dry rice. Phylogenetic clustering analyses demonstrate that these varieties probably belong to the tropical japonica variety (Figure 5). Of the 38 varieties analyzed, 37 form a highly supported monophyletic clade with the 3KRG tropical japonica accessions (Figure 5A). The short pairwise distances within this clade indicate relatively recent divergence and stand in sharp contract with the large pairwise distances to the other 3KRGP admixed and indica accessions (Supplementary Table S4). Principal component analysis confirms that the collected upland rice is similar to, but not identical with the tropical japonica accessions of the 3KRG project (Figure 5B). This is also reflected in the population structure analysis, which shows that the collected varieties share a higher proportion of genetic ancestry with the tropical japonica than with the indica and admixed accessions (Figure 5C). Only one collected sample stands out: Watralanti (nr. 6754), which means ‘water land’ and appears to be a wetland variety that bears more similarity to the 3KRGP indica (Figure 5A Group I) and particularly the admixed varieties Acorni and Ciwini (Figure 5A Group II) that were developed in Suriname in the 1970s (Supplementary Table S2). Watralanti is always planted near a creek or at the bottom on a slope as it needs moist soil. This rice probably descends from a wetland variety that was exchanged with East Indian rice farmers in coastal Suriname several decades ago.

Since our study area was divided in six collection sites, we checked for systematic differences in genetic diversity between sites. Pairwise genetic distances were on average not smaller (and even a bit higher) within the same geographic group (0.0137) then between geographic groups (0.0135) and closely resembled the population wide average genetic distance (0.0136). This shows that an even level of diversity of landraces is maintained in the different sampling locations that is not structured geographically, possibly because of gene flow. To test whether locations that are close together are more likely to exchange rice genetic diversity, we assessed genetic differentiation between populations along the North-South axis, following the Lawa/Maroni watershed that serves as the main highway for transportation of people and goods. In line with the low genetic distances among tropical japonicas, the fixation index (Fst) was found to be extremely low between sites and frequently less than 1%. This is exceedingly so between the island populations near Providence, Grand Santi and Mofina, which exhibit almost complete panmixia. Population differentiation is the highest (>2%) between the far northern areas of St. Laurent and the southern areas around Grand Santi and the Gonini river. This is consistent with a view of the Lawa/Maroni river as a transmission vector, along which rice is exchanged and diversity is actively maintained. Rather than a barrier against gene flow, the river is thus a place where gene pools meet and mix. This explains why the genetic differentiation between the opposite sides and countries is negligible, and why the legal border between French Guiana and Suriname does not form a natural border that leads to isolation.

#### 3.4.2 Improved varieties

It is noteworthy that our collection of landraces (with the exception of Watralanti), although morphologically heterogeneous, appears to forms a single large population without much genetic or geographic substructure. The cross-validation error estimate in ADMIXTURE was lowest at K=2 ancestral populations, with one of the ancestral populations accounting for most of the genetic makeup of the varieties (Supplementary Figure S3A). Only at K=4, we begin to differentiate groups within the Maroon varieties beyond the subspecies divide (Supplementary Figure S3B), but with the present data, this model is not supported. When we zoom in on the genetic variation among the 37 rainfed *O. sativa* collections, though, we do observe a clear separation between a group of genetically and morphologically more uniform improved varieties, that presumably entered Maroon agriculture some 70 years ago (Figure 5, Group III) and the larger and more diverse group of traditional varieties (Figure 5, Group IV). The internodal distance separating the improved clade is noticeably larger than the internodal distances among the traditional varieties (Supplementary Figure S4A). The narrow genetic base of this group is particularly apparent when we limit the two-dimensional scaling of genetic variation to the upland rice varieties (Supplementary Figure S4B). In this analysis, we see that most of the genetic space is occupied by the traditional landraces, whereas the improved varieties cluster relatively close together. To confirm that these modern varieties (Group III) are genetically less diverse than the traditional landraces (Group IV), we computed the average nucleotide diversity in 150 kb windows for both groups and compared them with a one-tailed Wilcoxon Rank Sum test. This showed that at 0.00011, the average divergence per window of Group III is significantly lower than that of Group IV at 0.00013 (p < 0.001).

This is of interest, because Group III includes the two most frequently cultivated varieties. The most popular one is known as ‘Mesti’ and grown by 29% of the farmers we interviewed. This term means ‘teacher’ in Ndyuka, but only one farmer remembered that this variety was named after a teacher who came to work at the first boarding school established in the region by the Moravian church on Stoelmanseiland, the island located in the confluence of the Tapanahoni and Lawa Rivers (Figure 1). As children from remote Maroon villages stayed on the island during the semester and there were few shops in the area, the teacher handed out rice to the mothers to grow on their own fields. The rice was subsequently supplied to feed the boarding school pupils. The white-husked Mesti rice quickly became popular and is still grown today. Farmers said it took longer to mature than most other varieties and compared its thin, elongated, slightly curved grains to the commercial ‘semi-super’ rice sold in shops. We could not trace the name of the teacher who introduced this variety, but it must have happened shortly after World War II, when the Moravians started their first mission center on Stoelmanseiland (Jabini, 2012). Vaillant (1948) did not encounter this variety in 1936, but a Maroon rice variety named ‘Mesiti’ was reported in the area in 1952 (Geijskes, 1955). We do not know the exact origin of Mesti rice.

Another popular and closely related variety is the ‘white Alekisoola’ or ‘white Sola’, grown by four farmers. It is considered as a ‘beautiful rice’, which is often sown earlier than other varieties, as it takes long to mature. It is planted on slopes and cannot tolerate drought. The rice does not break during dehusking and its grains are said to be ‘long like the commercial rice from Nickerie’. A drawback is that heavy rains during the harvest time cause the grains to become black and brittle. Varieties with this name were also reported by Geijskes (1955) and Baumgart et al. (1998) and identified as locally adapted farmers’ selections of Rexoro. This glabrous-hulled cultivar was developed in 1926 in Louisiana (Rutger and Mackill, 2001), introduced to Guyana in 1932 (Codd and Peterkin, 1933) and widely grown in coastal Suriname by1938. According to Stahel (1944), a bale of ‘Rexora seed padi’ was sent to the Saramaccan village Ganzee in 1936 and a few years later it was already grown by Ndyuka’s in the Maroni area. When Vaillant did his survey in 1936, it was not yet present there. The cultivation of Rexoro was discontinued in the 1960s because of a serious attack of Cercospora disease (Sanderson, 1962), but Alvarez et al. (2007) recently encountered it again on traditional farms in Cuba, decades after the introduction of modern varieties by the Cuban rice breeding program. The red-husked types of (Aleki-) Sola that we encountered have different morphological features than the white-husked types. They are sown and harvested together with the other rice varieties, with which they are genetically more closely related (Figure 5).

#### 3.4.3 Traditional varieties

Whereas the traditional Maroon varieties are cultivated less frequently, this is a larger group with higher genetic and phenotypic diversity (Figure 5, Group IV). Farmers often distinguish two types of a specific traditional variety, which are sown, harvested and stored separately: a ‘white’ one with straw-colored husks and a ‘red’ one, with orange-brown husks (Supplementary Table S4). Some women assured us that the only difference between these two types is the color, but our clustering analysis (Figure 5) shows that in no less than six varieties the white-husked types (of Onini, Sola, Alekisola, Alulu, Apiikutufutu, Abadagai and Bau anu) are not nearest neighbors with the red-husked types, but end up in different clades. In other cases, where varieties had the same name and the same husk color (such as Alulu, Weti alisi and Lebi Alisi), we found that they were phylogenetically separated. This shows that shared local names are not necessarily rooted in shared ancestry.

Several varieties with purple stems are named ‘Bau anu’ (blue hand), as the anthocyanins in the panicles stain people’s hands blue during the harvest. These varieties are not related and cluster phylogenetically with other varieties known under another local name. The blue sap is not seen as a negative trait; blue hand rice is said to be tasty, although it does not always give a good yield. One of the blue hand varieties (no DNA sequenced) is also known as ‘Ayengena’ (‘it does not hang’ in Ndyuka, after its somewhat erect panicles). In the early 1900s, Codd and Peterkin (1933) reported several varieties with purple pigmentation in Guyana, but we lack information on whether these are related to present-day Maroon varieties.

Six morphologically different varieties are named ‘Alulu (a bon)’, which literally means ‘it rolls (from the tree)’ or ‘it crumbles’ (Supplementary Table S4). The ripe seeds of this rice drop easily from the ear, so farmers cut the panicles just before they are fully mature and immediately place them in a bag to prevent the seeds from falling on the ground. The loss of seed shattering was a key event in the domestication of rice, as shattering caused a severe reduction of yield (Konishi et al., 2006). Maroon farmers, however, see the shattering as a positive trait, as it facilitates the threshing process. They praise this type of rice for its tasty, fat, white grains and ease of milling, although it is not tolerant to drought. The three ‘Alulu’ types that we sequenced cluster in different clades, so the shattering trait may have been reacquired several times independently or persisted from an ancestral gene pool.

Wintawaai (nr. 6809) is a variety that quickly lodges when there is much wind during the harvest season, after which the seeds shatter and rot. Lodging is generally seen as an undesirable trait of inferior landraces (Kashiwagi et al., 2005), but according to the single farmer who planted this variety, it grows well, is tasty and just needs to be cut before it is fully ripe.

One type of red-husked rice (nr. 6800 Lebi alisi) found near Providence, upper Maroni River, grouped together with the historic Wanica variety that was sequenced by Wang et al. (2018) in the 3000 Rice Genomes Project (Figure 5). This four-month variety was reportedly grown by Maroons in the 1930s in the interior of Suriname (Stahel, 1933). Two of our Maroon varieties (Aluku paansu and Abadagai) grouped close to one of the reference varieties from French Guiana (IRIS_313_7922, Figure 5), but no further information was provided on the origin of this accession.

#### 3.4.4 Ancestor varieties

In 1936, Maroons told Vaillant (1948) that according to their legend, rice came from Africa and was brought to Suriname by a female ancestor who concealed the grains in her hair (Carney, 2009). We found this legend still to be known in St. Laurent today, and videotaped a demonstration of how a Maroon woman braids hands full of rice grains invisibly into her daughter’s hair (Supplementary Material 1, video 6). She explained that besides rice, other crop seeds and cassava cuttings were also smuggled this way, both in Africa when women were enslaved and in Suriname when Maroons fled to into the forest. Moreover, we encountered two rice varieties in Bigiston that still carry the names of the enslaved women who brought this rice to the interior. The variety ‘Milly’ (nr. 6760) is named after an escaped female ancestor ‘who introduced this rice to the Maroni River in times of slavery when people did not yet live in the forest […] when they had not yet created villages in the woods’. Another variety, growing in the same field, is named ‘Sapali’ (nr. 6761), after an enslaved woman who, after she fled from the plantation, crossed a savanna where she encountered rice plants, from which she took ripe seeds that she shared with other runaways. Although Sapali is appreciated for its ‘fat, white and clean’ grains, the scabrous leaves scratch your skin and the seeds are difficult to dehusk by hand. According to Ndyukas along the Cottica River interviewed around 1961 by anthropologist Köbben (1968), their ancestors had suffered from hunger after their escape from the plantations, until they were joined by a female runaway named Ma Sapá, who had hidden rice kernels in her hair. The current Sapali rice likely descends from an early 18^th^ century variety that has been crucial to the survival of the Ndyuka Maroons.

#### 3.4.5 Obsolete and weedy rice varieties

The exact number of Maroon rice varieties is probably higher than 50, as we missed some varieties that had been planted in 2017, but had already been consumed, such as the red and white types of the Saramaccan varieties 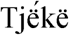 muyee and Agbonso. Farmers assured us that many more varieties could be found in traditional villages along the remote Tapanahoni and upper Suriname Rivers. Some women told us about varieties planted by their mothers that are not sown anymore nowadays, because of their long, itching awns or scabrous leaves, which are seen as a nuisance. Examples are ‘Saapa’ (probably referring to the Sapali variety) and rice types aptly described as ‘pubic hair’ after their long, curled black awns. The latter varieties (nrs. 6758, 6767) are still grown by some farmers, who said that ‘they look beautiful, give a good yield and we enjoy planting it’. One farmer said she was ‘surprised that people still grow this old-fashioned rice’, when we showed her our collections of awned, shattering and lodging varieties. Two of the tropical japonica varieties that were sequenced by Wang et al. (2018) go by a similar name (Pinde gogowierie). Like Wanica, many of the 3KRGP accessions were collected and deposited in seed banks several decades ago, which means they do not necessarily represent the currently used gene stock.

In addition, there was much morphological variation in the types that carried the generic names ‘weti alisi’ and ‘lebi alisi’ (white resp. red rice), but we were not able to extract DNA from every single accession. On the upper Maroni islands near the village of Providence (Figure 1), we collected two samples of ‘white rice’ (nrs. 6799, 6801) that turned out to be mixtures of grains with pubescent and glabrous husks, dark and uncolored tips, white and red pericarp (Supplementary Table S4). The red-seed rice (nrs. 6801A and 6799B) is sometimes called ‘lebi wata’, referring to the red color of the water when it is cooked, but it is not stored separately from varieties with white pericarp. Apparently, the samples were infested with ‘weedy’ or ‘red’ rice, sometimes indicated as *Oryza sativa* f. *spontanea* Roshev., although this name is considered to be a synonym of *O. sativa* (The Plant List, 2013). In the 1920s, the ‘predominance of a particularly undesirable type of wild rice with red grains’ among traditional landraces was one of the reasons for the selection of pure lines for commercial rice cultivation in Guyana (Codd and Peterkin 1933). Weedy rice is still a serious agricultural problem in directly sown rice worldwide, and various types exist (with or without awns and with different husk colors), often mimicking the phenotype of the desired variety (Londo and Schaal, 2007). These red-seeded types can arise when Asian rice hybridizes with its wild ancestor, *Oryza rufipogon* Griff., creating offspring that shatter quickly, have long, barbed awns, increased competitive ability, dormant seeds and a red pericarp. Weedy rice also develops when domesticated rice is abandoned and turns feral (Ellstrand et al., 2003). The weedy rice types we encountered probably belong to the ‘strawhull type’ (Gealy, 2005; Londo and Schaal, 2007), as they lack awns, have straw-colored husks and red pericarp. Although of Asian origin (Stein et al., 2018), *O. rufipogon* occurs in the wild in the Guianas (Judziewicz, 1990). The herbarium label of a specimen from 1978 (P. Grenand nr. 1633, CAY) indicated that Palikur Indians in French Guiana consumed this wild rice in the past, although the awns were difficult to remove. Future genetic analyses can prove whether the weedy rice we encountered represent hybrids between traditional varieties and wild rice or escaped rice gone feral.

The preceding sections show a remarkable array of modern, improved, traditional and ancient landraces, ranging from commonly used to almost disused. Phenotypic diversity and vernacular names are distributed rather homogeneously both in phylogenetic (Figure 5) and in geographic space (Figure 1).

## 4 Discussion

### 4.1 More hidden rice diversity

Since the 1936 expedition by Vaillant, our study is the first inventory of Ndyuka Maroon rice that is supported by herbarium vouchers, living seeds safeguarded at a germplasm institute and genetic analysis. We rediscovered only seven of the ± 30 Ndyuka rice names documented by Vaillant (1948), five of the 21 Aluku names listed by Fleury (2016), four of the 25 Saramaccan names of Baumgart et al. (1998) and five of the 23 Ndyuka names published by Geijskes (1955). On the other hand, 15 of the local rice names that we documented had never been reported before. We cannot tell which of the previously documented varieties have been lost in the past 80 years, as our inventory covered only a small part of the Maroni and Lawa Rivers, rice names are subject to change, and Vaillant’s herbarium vouchers are not digitally available. Price and Price (2017) suggested a substantial decline in the number of Maroon rice varieties over the past decades, but no collections or name lists are available to prove this assumption. Further morphological and genetic comparison of historic rice collections (such as Vaillant’s vouchers at the Muséum National d’Histoire Naturelle in Paris) with recent samples and additional field collecting are needed to provide a more complete overview of the extraordinary rice diversity in this area, its dynamics in space and time and the survival of ancient landraces.

Along the Maroni-Lawa watershed, farmers cultivate a large number of rice varieties, each with its own morphological and agronomical characteristics. This phenotypic diversity, ranging from ancient African landraces (*O. glaberrima*) to relatively recent introductions (US cultivars of *O. sativa* subsp. *japonica* and a Hindostani wetland variety) reflects the history of the human migration in the region and their contacts with outsiders. This diversity also reflects the multiple objectives of rice cultivation for farming households: it is simultaneously a food production strategy, a social and ritual asset. Foreign varieties introduced at different times in the history of the region have contributed to increase the phenotypic diversity. The fact that the phenotypic diversity described in this paper is apparent at all collection sites, suggests a certain level of gene flow between the populations along the river, serving as a pathway for seed exchange. This is concordant with the lack of geographic and genetic structure observed in our population genomic analyses (Figure 5).

Exchange of seed material within ethno-linguistic groups generally occurs at higher frequencies than between different groups, as seed exchanges are based on cultural preferences (Labeyrie et al., 2014). Ndyuka farmers hardly trade rice with other Maroon groups, and they ensured us that ‘Saramaccans had different types of rice’. This is in line with recent sorghum studies showing that the spatial distribution of landrace names and genetic diversity of traditional crops are significantly correlated with ethnolinguistic partition (Westengen et al., 2014; Labeyrie et al. 2016). We therefore think that the diversity of landraces that we encountered represents only the tip of the iceberg of hidden rice diversity in the Guianas, and that similar ethnobotanical surveys among other Maroon groups will reveal additional varieties not covered in this study. To date, no voucher material exists for rice varieties of the Paramaccan Maroons who live along the middle Maroni River or the Aluku Maroons along the upper Lawa. Only a few collections have been made among Saramaccan Maroons (Baumgart et al., 1998; Van Andel et al., 2016b), but no research has been done on rice grown by the Matawai and Kwinti Maroons in central Suriname. Apart from a paper on rice cultivation by Hmong migrants from Laos in French Guiana (Salaul □n, 1999), no studies or herbarium vouchers exist of traditional rice varieties currently grown by descendants of Asian migrants in the Guianas. This study is therefore the first step towards obtaining a full picture of the landraces that are still in cultivation by different ethnolinguistic groups in the Guianas.

### 4.2 Where did the Maroon rice varieties come from?

Several rice varieties collected during this study possessed traits that are characteristic to traditional landraces, such as tall stems, awns, scabrous leaves, pubescent husks, shattering seeds, small grains and a proneness to lodging. Awns and rough hairs help to protect seeds from predation by birds and mammals, and aid seed dispersal by clinging to animal fur. However, awns and scabrous plant parts also hinder mechanical seed processing and storage, so modern rice cultivars have been selected to be glabrous, awnless or short-awned (Rutger and Mackill, 2001; Hua et al. 2015). Greater plant height makes rice more susceptible to lodging, which is considered a negative trait for machine-harvested rice, so modern cultivars have been selected for short and sturdy stems (Nascente and Kromocardi, 2017). However, lodging does not pose a problem if panicles are cut by hand (Kashiwagi et al., 2005). The development of non-shattering cultivars has also been a strong focus of rice breeding programmes, although modern indica cultivars are still more prone to exhibit seed shattering than japonica cultivars (Konishi et al., 2006). Typically, all Maroon accessions we sequenced are tropical japonicas, and several of them shatter quickly, just like our sample of *Oryza glaberrima*.

In 1927, the national rice breeding programme in Guyana started to develop pure lines from locally grown rice landraces by removing volunteer plants and selecting non-shattering, awnless strains with a high yield, strong straw, uniform plant height and white grains to facilitate mechanical threshing and milling. These new cultivars were handed out to farmers to ensure the ‘elimination of traditional varieties with undesirable characteristics’ (Codd and Peterkin, 1933:2-3). In the 1940s, through World War II, the Dutch started investments in commercial rice breeding in Suriname. They imported the pure lines from Guyana and improved cultivars from Java, Indonesia and the US for their field trials, but also tested out some varieties that were probably of Maroon origin, like Wanica. After the war, modern wetland cultivars were developed for mechanized rice cultivation in the coastal polders (Maat and Van Andel, 2018).

It is likely that some of the varieties we collected descend from these pre-1927 landraces, although the manifold origins of the Maroon rice diversity are difficult to trace. Europeans introduced the crop in the Guianas in the early colonial period, but the exact routes and motivations remain unclear (Maat and Van Andel, 2018). Plantation holders imported hulled rice from Louisiana and Carolina as provision for the slaves (Codd and Peterkin, 1933; Rolander, 2008). Jewish planters that were expelled from Brazil may have introduced rice to Suriname when they migrated there around 1667 (Young and Angier, 2010). Slave traders bought stocks of rice from merchants along the West African coast to feed their captives during the Middle Passage (Carney 2009; Mouser et al., 2015; van Andel et al., 2016a). *Oryza sativa* was introduced to West Africa in the 16th century, before the onset of the transatlantic slave trade, and adopted by peoples along the upper Guinea Coast who had previous experience growing the local African species (Linares, 2002). Vaillant (1948) suggested that most of the Maroon rice diversity had an African origin. However, Saramaccan rice farmers interviewed by Price (1993) said that while in their youth (in the 1920s), there were only a handful rice varieties (black rice was one of them), diversity increased substantially in the 1960s when Maroon men took up wage labor in coastal Suriname and returned with planting stock exchanged with East Indian and Javanese farmers. Therefore, part of the Maroon rice diversity may have originated from landraces introduced by Asian contract laborers.

Recently, the genome of a single accession of black rice from Suriname was compared to 109 accessions of *O. glaberrima* collected across West Africa. A strong similarity was established with a landrace grown in western Ivory Coast (Van Andel et al., 2016a), which shows how genomics can reveal unwritten migration histories of crop varieties. Applying comparative phylogenomics to historic rice collections from the Guianas in museums and gene banks and currently cultivated Maroon landraces, and comparing them with traditional varieties and cultivars from Africa, Asia and the US can shed a light on the geographic origins of Maroon lineages and reveal patterns of crop migration and adaptation that have remained hidden for centuries. Special attention should be paid to varieties that Maroons consider to have been introduced by enslaved ancestors, such as Milly and Sapali for the Ndyuka and Paánza rice for the Saramaccans (Price, 2002), of which one accession is stored at the SNRI/ADRON seed bank.

### 4.3 Future of Maroon rice

The Maroon forest garden represents the classical slash-and-burn agroecosystem (Kleinman et al., 1995), in which rice is sown once on a freshly burned field and intercropped with cassava, maize, bananas and vegetables. Without the use of fertilizers or pesticides, the field is left fallow after one season, after which only volunteer rice plants and perennial crops are harvested. Farmers spread their risk by planting several distinct rice varieties on one field. Early and late maturing types are sown in succession to even out labor during harvest time. In this way there is a higher chance that at least one variety will thrive, as the weather may be unpredictable, and wet and dry seasons do not always start and end at the same time over the years. Cropping patterns that mimic ecological complexity are most effective at soil conservation and promoting sustainability, as long as rotation periods are long and human population pressure is low (Kleinman et al., 1995).

Governments and agricultural development programs generally promote continuous cropping systems over shifting cultivation (Kleinman et al., 1995) and Suriname is no exception to this phenomenon (Fleskens and Jorritsma, 2010). After visiting the Maroon rice fields, Geijskes (1955:136) claimed that yields were not always sufficient and ‘due to the wasteful land use, the agriculture of the Bush negroes has become a matter which urgently needs the attention of the government’. Tropical plant breeders Budelman and Ketelaars (1974:17) advised that dryland rice cultivation in the interior should be terminated, ‘as this is a difficult crop for permanent cultivation on these grounds’ and recommended oil palm plantations as a more suitable cash crop. In 1992, the French government made violent attempts to force the Maroon refugees from the Surinamese civil war to leave the area and sprayed pesticides on their rice fields (Léobal, 2016). Maroon farmers around St. Laurent, illegal immigrants and legal French citizens alike, considered the ongoing harassment by the French police as a serious problem affecting their agricultural practices.

As a remedy for short fallow periods and rice yields that did not meet the local demand, caused by population pressure, Baumgart et al. (1998) suggested to intensify Maroon rice cultivation by purifying traditional varieties, introducing improved upland cultivars, fertilizers, herbicides and mechanization. However, in their study on soil fertility on Maroon fields, Fleskens and Jorritsma (2010) discovered that due to migration to the city, the issue of human pressure on the land was rarely raised by Maroon farmers. In a recent experiment, Nascente and Kromocardi (2017) tried to increase yields by adding NPK fertilizer, herbicide and fungicide to three traditional Saramaccan landraces and three improved Brazilian upland cultivars. Only one of the Maroon varieties responded positively to this treatment, after which they recommended to switch to Brazilian cultivars to double the current average of upland rice production and improve food security among the Maroon population. They acknowledged, however, that the ‘very limited financial capital and low education level’ of the Maroons limited their access to modern technologies and that the introduction of high-yield rice cultivars required subsidies for farmers to access fertilizers and pesticides (Nascente and Kromocardi, 2017:192).

In addition to the governmental pressure, the upcoming evangelical churches in the Maroon territories are rigidly opposed to Afro-religious practices, which has led to frictions within communities (Van Stipriaan, 2015). According to Richard Price (pers. comm. 3 May 2018), converted Christians are increasingly discouraged to visit funerals in remote Maroon villages, as these are regarded as places of witchcraft. Since funerals are core events at which landraces are exchanged, processed and consumed, this phenomenon could affect the survival of Maroon rice diversity. Traditional crop varieties can quickly disappear if not sown every few years. In Grand Santi, however, several members of evangelical churches still cultivate rice, including black rice, for home consumption and sale.

In spite of half a century of policy recommendations to change or abandon their traditional agriculture, Maroon rice farming has shown a remarkable resilience. Commercial rice has entered their diet decades ago (Geijskes, 1955; Bilby et al., 1989), but did not replace their own landraces. New waves of Ndyuka Maroons arrived in French Guiana as refugees during the Surinamese civil war in the 1980s, which led to a rapid urbanization of the region around St. Laurent du Maroni and the abandonment of agriculture (Fleskens and Jorritsma, 2010; Léobal, 2016). This provided more farmland for those that stayed in their forest villages and created a market for homegrown rice in St. Laurent. Mechanical rice mills, often donated in development projects (e.g., Rosebel Gold Mines, 2017; De Ware Tijd, 2018), can lead to malnutrition and thiamine deficiency as they remove the nutritious bran and germ of the rice (Lanska, 2010; Ryan, 2011). But Maroon farmers are well aware of this, and have not given up the traditional dehusking methods with mortar and pestle.

The Maroons, however, do not have to remain solely responsible for protecting their cultural heritage. The fascinating history of Maroon rice, its unique diversity, dynamic character, distinctive taste and strong gender dimension (which is reflected in varieties known as ‘dancing woman’, ‘beautiful woman’ or the names of female ancestors) offer opportunities for broader culinary marketing and ‘agro-ecotourism’ (see Maxted et al., 2002). The general public in the Guianas is hardly aware of Maroon rice. Not a single restaurant in Cayenne or Paramaribo sells Maroon rice dishes. Given the increased interest by global consumers in traditional food products (Ardenghi et al, 2018), a greater awareness of the unique Maroon rice varieties could stimulate their conservation in the face of increased urbanization and outside pressures. An appreciation of the role of Maroons as custodians of rice diversity would benefit not just local food security, but also safeguard a precious global resource.

## Supplementary Material

The Supplementary Material for this article can be found online at:

## Author Contributions statement

TA, HM, and HB conceived and designed the study, DHL and TP gave input on the study design, TP and TA organized fieldwork, JT provided the data collection format, germplasm storage and phenotyping data, TA and AB carried out the fieldwork, AB, MV, VM and HB carried out the genetic research. All authors reviewed and approved the final manuscript.

## Acknowledgements

We are grateful to the staff of SNRI/ADRON in Nickerie, the Herbier IRD in Cayenne, and to Frédéric Blanchard (Collectivité Territoriale de Guyane, Cayenne), Brian Mawdo, Manon Plasschaert, Beatrice Rostand and Lani Pesna for facilitating our fieldwork. We thank Frans Afri, Stieven Scheinemann and Edith Adjako for their work as translators. We are indebted to rice farmers Maneska Manu, Martha Afonsoewa, Cynthia Asoiti, Elise Sandiana, Nesia Atanso, Christine Audo, Samantha Dansman, Christine Feno, Comina Eeswijk, Eleni Galimo, Sonia Sini, Ajadie Abetau, Ifna Asaida, Marceline and Vanessa Kodeli, Jomea Nyanfai, Agnes Awenkina, Sylvie and Maloe Deel, Leonie Altret and Paisie Mbola for sharing their knowledge and their rice varieties with us. We thank Sally Price for sharing her unpublished data on Saramaccan rice varieties. Linh Nguyen Nhat helped us with the lab work at DNA lab of the Natural History Museum of the University of Oslo (UiO), Norway. This work was performed on the Abel Cluster, owned by the University of Oslo and the Norwegian metacenter for High Performance Computing (NOTUR), and operated by the Department for Research Computing at USIT, the University of Oslo IT-department, http://www.hpc.uio.no/.

---

**Figure 4.** *Oryza glaberrima* seeds (nr. 6782), showing the black husks and straight awns. Picture by Tinde van Andel.

**Figure 5.** Position of Maroon landraces within the Asian rice gene pool. **A.** Approximately Maximum Likelihood (ML) tree of 39,595 homozygous SNPs. Branch lengths are not shown. Branches with low support values (<0.70) are collapsed. Support values of the remaining branches are indicated on a scale from 0.70 (red) through 0.85 (yellow) to 1.00 (green). **B.** Principal component analysis of 64,313 unlinked SNPs. **C.** ADMIXTURE analysis of 64,313 unlinked SNPs at K=2 ancestral populations.

**Figure 6.** Two shattering varieties of Maroon rice named ‘Alulu’ with husked (left) and dehusked (right) grains. **A.** ‘White type’ with straw-coloured, pubescent, black-tipped husks (nr. 6772A) and **B.** ‘Red type’, with orange-brown, glabrous, black-tipped husks (nr. 6819). Pictures by Alice Bertin.

## References

Alexander DH, Lange K. Enhancements to the ADMIXTURE algorithm for individual ancestry estimation. BMC Bioinformatics (2011) 12: 246. https://doi.org/10.1186/1471-2105-12-246

Alvarez A, Fuentes JL, Puldón V, Gómez PJ, Mora L, Duque MC et al. Genetic diversity analysis of Cuban traditional rice (Oryza sativa L.) varieties based on microsatellite markers. Genet Mol Biol (2007) 30:1109-17. http://dx.doi.org/10.1590/S1415-47572007000600014

Ardenghi NMG, Rossi G, Guzzon F. Back to beaked: Zea mays subsp. mays Rostrata Group in northern Italy, refugia and revival of open-pollinated maize landraces in an intensive cropping system. PeerJ (2018) 6:e5123. doi: 10.7717/peerj.5123

Baumgart IR, HilleRisLambers D, Khodabaks MR, Wildschut J. Visit to rice growing sites on the upper Suriname river between Nieuw Aurora and Abenaston. Nickerie: ADRON (1998). 6 p.

Bergey CM. vcf-tab-to-fasta. (2012). http://code.google.com/p/vcf-tab-to-fasta

Bilby K, Delpech B, Fleury M, Vernon D. L’alimentation des noirs Marrons du Maroni: vocabulaire, pratiques, representations. Cayenne: ORSTOM (1989). 393 p.

Bolger AM, Lohse M, Usadel B. Trimmomatic: a flexible trimmer for Illumina sequence data. Bioinformatics (2014) 30:2114–20. https://doi.org/10.1093/bioinformatics/btu170

Budelman A, Ketelaars JJMH. Een studie van het traditionele landbouwsysteem onder de boslandcreolen. Paramaribo: CELOS (1974). 148 p.

Carney JA. Black rice: the African origins of rice cultivation in the Americas. Boston: Harvard University Press (2009). 256 p.

Clément G, Dance D, Cavana JF. Quel avenir pour la filière rizicole de Mana? Montpellier: CIRAD (2011). 72 p.

Codd LEW, Peterkin EM. Rice in British Guiana, 1927-1932. Georgetown: British Guiana Department of Agriculture Rice bulletin (1933) 1:1–38.

Danecek P, Auton A, Abecasis G, Albers CA, Banks E, DePristo MA, et al. The variant call format and VCFtools. Bioinformatics (2011) 27:2156–58. https://doi.org/10.1093/bioinformatics/btr330

De Ware Tijd. 2018. Conservation International doneert aan Matawai. http://dwtonline.com/laatste-nieuws/2018/04/18/conservation-international-doneert-aan-matawai/

Delnatte C, Meyer JY. Plant introduction, naturalization, and invasion in French Guiana (South America). Biol Invasions (2012) 14:915–27. https://doi.org/10.1007/s10530-011-0129-1

Ellstrand NC, Prentice HC, Hancock JF. Gene flow and introgression from domesticated plants into their wild relatives. Annu Rev Ecol Syst (2003) 30:539–63. https://doi.org/10.1146/annurev.ecolsys.30.1.539

Fleskens L, Jorritsma F. A behavioral change perspective of Maroon soil fertility management in traditional shifting cultivation in Suriname. Hum Ecol (2010) 38:217–36. https://doi.org/10.1007/s10745-010-9307-5

Fleury M. Racines alimentaires: L’alimentation des Noirs marrons en Guyane française. Hommes et Plantes (2012) 83:9–17.

Fleury M. Agriculture itinérante sur brûlis (AIB) et plantes cultivées sur le haut Maroni: étude comparée chez les Aluku et les Wayana en Guyane française. Bol Mus Para Emílio Goeldi (2016) 11:431–65. http://dx.doi.org/10.1590/1981.81222016000200006

Gealy DR. “Gene movement between rice (*Oryza sativa*) and weedy rice (*Oryza sativa*- a US temperate rice perspective,”. In: Gressel J, editor. Crop Ferality and Volunteerism. Boca Raton, FL: CRC Press (2005). p. 323–54.

Geijskes DC. De landbouw bij de bosnegers van de Marowijne. De West-Indische Gids (1955) 55:135–53.

Heemskerk M. Gender and gold mining: the case of the Maroons of Suriname. East Lansing, MI: Women in International Development, Michigan State University (2000). 33 p.

Heemskerk M. Scenarios in anthropology: reflections on possible futures of the Suriname Maroons. Futures (2003) 35:931–49. https://doi.org/10.1016/S0016-3287(03)00050-8

Heinemann RJB, Fagundes PL, Pinto EA, Penteado MVC, Lanfer-Marquez UM. Comparative study of nutrient composition of commercial brown, parboiled and milled rice from Brazil. J Food Compost Anal (2005) 18:287–96. https://doi.org/10.1016/j.jfca.2004.07.005

Hua L, Wang DR, Tan L, Fu Y, Liu F, Xiao L et al. LABA1, a domestication gene associated with long, barbed awns in wild rice. Plant Cell (2015) 27:1875–88. https://doi.org/10.1105/tpc.15.00260

Hurault J. La vie matérielle des noirs réfugiés Boni et des Indiens Wayana du Haut-Maroni (Guyane française). Paris: ORSTOM (1965). 145 p.

Jabini FS. Christianity in Suriname. Carlisle: Langham Partnership (2012). 416 p.

Jackson MT, Lettington RJL. Conservation and use of rice germplasm: an evolving paradigm under the international treaty on plant genetic resources for food and agriculture. Rome: FAO (2002). http://www.fao.org/docrep/006/Y4751E/y4751e07.htm

Johnson MG, Pokorny L, Dodsworth S, Botigué LR, Cowan RS, Devault A, et al. A universal probe set for targeted sequencing of 353 nuclear genes from any flowering plant designed using k-Medoids clustering. Syst Biol (2018). https://doi.org/10.1093/sysbio/syy086

Judziewicz EJ. “Poaceae (Gramineae),”. In: Gorts-van Rijn A, editor. Flora of the Guianas, Series A, Phanerogams 8. Koenigstein: Koeltz Scientific Books (1990). p 1-727.

Kashiwagi T, Sasaki H, Ishimaru K. Factors responsible for decreasing sturdiness of the lower part in lodging of rice (Oryza sativa L.). Plant Prod Sci (2005) 8:166–72. https://doi.org/10.1626/pps.8.166

Kleinman PJA, Pimentel D, Bryant RB. The ecological sustainability of slash-and-burn agriculture. Agric Ecosyst Environ (1995) 52:235–49. https://doi.org/10.1016/0167-8809(94)00531-I

Köbben AJF. Continuity in change-Cottica Djuka society as a changing system. Bijdr Taal Land Volkenkd (1968) 124: 56–90.

Konishi S, Izawa T, Lin SY, Ebana K, Fukuta Y, Sasaki T et al. An SNP caused loss of seed shattering during rice domestication. Science (2006) 312:1392–6. doi: 10.1126/science.1126410

Korneliussen TS, Albrechtsen A, and Nielsen R. ANGSD: analysis of next generation sequencing data. BMC Bioinformatics (2014) 15:356. https://doi.org/10.1186/s12859-014-0356-4

Kumar S, Stecher G, Tamura K. MEGA7: Molecular Evolutionary Genetics Analysis version 7.0 for bigger datasets. Mol Biol Evol (2016) 33:1870–74. https://doi.org/10.1093/molbev/msw054

Labeyrie V, Deu M, Barnaud A, Calatayud C, Buiron M, Wambugu P et al. Influence of ethnolinguistic diversity on the sorghum genetic patterns in subsistence farming systems in Eastern Kenya. PLoS One (2014) 9:e92178. https://doi.org/10.1371/journal.pone.0092178

Lanska DJ. Historical aspects of the major neurological vitamin deficiency disorders: the watersoluble B vitamins. Handb Clin Neurol (2010) 95:445–76. doi: 10.1016/S0072-9752(08)02129-5

Lemmon EM, Lemmon AR. High-throughput genomic data in systematics and phylogenetics. Annu Rev Ecol Evol Syst (2013) 44:99–121. https://doi.org/10.1146/annurev-ecolsys-110512-135822

Léobal C. “From “primitives” to “refugees”: French Guianese categorizations of Maroons in the aftermath of Surinamese civil war,”. In: Hassankhan MS, Roopnarine L, White C, Mahase R, editors. Legacy of Slavery and Indentured Labour: Historical and Contemporary Issues in Suriname and the Caribbean. New York: Routledge (2016). p. 213–30.

Li H, Durbin R. Fast and accurate short read alignment with Burrows–Wheeler transform. Bioinformatics (2009) 25:1754–60. https://doi.org/10.1093/bioinformatics/btp324

Li H, Handsaker B, Wysoker A, Fennell T, Ruan J, Homer N. The Sequence Alignment/Map format and SAMtools. Bioinformatics (2009) 25: 2078–79. https://doi.org/10.1093/bioinformatics/btp352

Linares OF. African rice (Oryza glaberrima): History and future potential. PNAS (2002) 99:16360–5. https://doi.org/10.1073/pnas.252604599

Londo JP, Schaal BA. Origins and population genetics of weedy red rice in the USA. Mol Ecol (2007) 16:4523–35. https://doi.org/10.1111/j.1365-294X.2007.03489.x

Maat H, Van Andel TR. The history of the rice gene pool in Suriname: circulations of rice and people from the eighteenth century until late twentieth century. Hist Agr (2018) 75:69–91. http://dx.doi.org/10.26882/histagrar.075e04m

Mather KA, Caicedo A, Polato N, Olsen KM, McCouch S, Purugganan MD. The extent of linkage disequilibrium in rice (Oryza sativa L.). Genetics (2007) 177:2223–32. https://doi.org/10.1534/genetics.107.079616

Maxted N, Guarino L, Myer L, Chiwona EA. Towards a methodology for on-farm conservation of plant genetic resources. Genet Resour Crop Evol (2002) 49:31–46. https://doi.org/10.1023/A:101389640

Meyer M, Kircher M. Illumina Sequencing Library Preparation for highly multiplexed target capture and sequencing. Cold Spring Harb Protoc (2010) 6:pdb.prot5448. doi:10.1101/pdb.prot5448

Mouser BL, Nuijten E, Okry F, Richards, P. “Red and white rice in the vicinity of Sierra Leone: Linked histories of slavery, emancipation and seed selection,”. In: Bray F, Coclanis PA, Fields-Black EL, Schäfer D, editors. Rice: Global Networks and New Histories. New York: Cambridge University Press (2015). p. 138-62.

Nascente AS, Kromocardi R. Genotype selection and addition of fertilizer increases grain yield in upland rice in Suriname. Acta Amazon (2017) 47:185–94. http://dx.doi.org/10.1590/1809-4392201603374

Perales HR, Benz BF, Brush SB. Maize diversity and ethnolinguistic diversity in Chiapas, Mexico. PNAS (2005) 102:949–54. https://doi.org/10.1073/pnas.0408701102

Portères R. Présence ancienne d’une variété cultivée d’*Oryza glaberrima* St. en Guyane Française. J Agric Trop Bot Appl (1955) 2:680.

Price R. Subsistence on the plantation periphery: crops, cooking, and labour among eighteenth century Suriname Maroons. Slavery Abol (1991) 12: 107–27.

Price R. First-time: The historical vision of an African American people. Chicago, IL: University of Chicago Press (2002). 208 p.

Price R. Rainforest warriors: human rights on trial. Philadelphia, PA: University of Pennsylvania Press (2012). 280 p.

Price R. The Maroon population explosion: Suriname and Guyane. NWIG (2013) 87:323–27.

Price S. Co-wives and calabashes. Ann Arbor, MI: University of Michigan Press (1993). 264 p.

Price R, Price S. Saamaka dreaming. Durham: Duke University Press (2017). 272 p.

Price MN, Dehal PS, Arkin AP. FastTree 2: Approximately Maximum–Likelihood trees for large alignments. PLoS ONE (2010) 5:e9490. doi:10.1371/journal.pone.0009490.

Purcell S, Neale B, Todd-Brown K, Thomas L, Ferreira MA, Bender D, et al. PLINK: a tool set for whole-genome association and population-based linkage analyses. Am J Hum Genet (2007) 81:559–75. https://doi.org/10.1086/519795

Reijers M. African heritage in Maroon agriculture: multiple uses of Old World crops among Aucans and Saramaccans. Wageningen: Master thesis, Wageningen University (2014). 65 p.

Rolander D. “Daniel Rolander’s Journal (1754-1756), The Suriname Journal: composed during an exotic journey,”. (translated from Latin by Dobreff J). In: Hansen L, editor. The Linnaeus Apostles: Global Science & Adventure (2008) 3:1215–1576.

Rosebel Gold Mines. De bloeiende toekomst van Brokopondo. Het district, de gemeenschappen en hun inwoners. Paramaribo: VACO Publishers (2017). 86 p.

Rutger JN, Mackill DJ. “Application of Mendelian genetics in rice breeding,”. In: Khush GS, Brar DS, Hardy B, editors. Rice Genetics IV. Enfield, NH: Science Publishers (2001). p. 27–38.

Ryan EP. Bioactive food components and health properties of rice bran. J Am Vet Med Assoc (2011) 238:593–600. https://doi.org/10.2460/javma.238.5.593

Salauln P. Le système de production agricole Hmong à Saull (Guyane franclaise): modalités de pérennisation. JATBA (1999) 41:251–79. https://doi.org/10.3406/jatba.1999.3720

Sanchez PL, Wing RA, Brar DS. “The wild relative of rice: genomes and genomics,”. In: Zhang Q, Wing RA, editors. Genetics and genomics of rice, plant genetics and genomics: crops and models 5. New York: Springer (2013). p. 9–25. doi: 10.1007/978-1-4614-7903-1_2

Sanderson AG. The agricultural economy of Suriname (Dutch Guiana). Washington DC: US Department of Agriculture, Economic Research Service (1962). 30 p.

Stahel G. De rijstcultuur in Suriname. Paramaribo: Departement Landbouwproefstation in Suriname. Bulletin 2 (1933). 41 p.

Stahel G. De Nuttige Planten van Suriname. Paramaribo: Departement Landbouwproefstation in Suriname. Bulletin 59 (1944). 239 p.

Stein JC, Yu Y, Copetti D, Zwick DJ, Zhang L et al. Genomes of 13 domesticated and wild rice relatives highlight genetic conservation, turnover and innovation across the genus Oryza. Nature Genetics (2018) 50: 285–96. https://doi.org/10.1038/s41588-018-0040-0

Tareau MA, Palisse M, Odonne G. As vivid as a weed: medicinal and cosmetic plant uses amongst the urban youth in French Guiana. J Ethnopharmacol (2017) 203:200–13. https://doi.org/10.1016/j.jep.2017.03.031

The Plant List. Version 1.1 (2013). http://www.theplantlist.org/

Vaillant M. Milieu cultural et classification des variétés de riz des Guyanes française et hollandaise. Rev Int Bot Appl d’Agriculture Trop (1948) 28:520–29. https://doi.org/10.3406/jatba.1948.6700

Van Andel TR. African rice (Oryza glaberrima Steud.): Lost crop of the enslaved Africans discovered in Suriname. Econ Bot (2010) 64:1–10. https://doi.org/10.1007/s12231-010-9111-6

Van Andel T, Maas P, Dobreff J. Ethnobotanical notes from Daniel Rolander’s Diarium Surinamicum (1754-1756): are these plants still used in Suriname today? Taxon (2012) 61:852– 63. https://doi.org/10.1002/tax.614010

Van Andel TR; Meyer RS, Aflitos SA, Carney JA, Veltman M, Copetti D et al. Tracing ancestor rice of Suriname Maroons back to its African origin. Nat Plants (2016a) 2:10. https://doi.org/10.1038/nplants.2016.149

Van Andel TR, Van der Velden A, Reijers M. The ‘Botanical gardens of the Dispossessed’ revisited: richness and significance of Old World Crops grown by Suriname Maroons. Genet Resour Crop Evol (2016b) 63:695–710. https://doi.org/10.1007/s10722-015-0277-8

Van Stipriaan A. “Maroons and the communications revolution in Suriname’s interior,”. In: Carlin E, Léglise I, Migge B, Tjon Sie Fat P, editors. In and out of Suriname: language, mobility and identity. Leiden: Brill (2015). p 139–63.

Wang, W, Mauleon R, Hu Z, Chebotarov D, Tai S, Wu Z. Genomic variation in 3,010 diverse accessions of Asian cultivated rice. Nature (2018) 557:43–9. https://doi.org/10.1038/s41586-018-0063-9

Weir B, Cockerham C. Estimating F-Statistics for the analysis of population structure. Evolution (1984) 38:1358–70. https://doi.org/10.1111/j.1558-5646.1984.tb05657.x

Westengen OT, Okongo MA, Onek L, Berg T, Upadhyaya H, Birkeland S et al. Ethnolinguistic structuring of sorghum genetic diversity in Africa and the role of local seed systems. PNAS (2014) 111:14100–5. https://doi.org/10.1073/pnas.1401646111

World Bank Group. Climate change knowledge portal for development practitioners and policy makers. Data for Suriname 1991-2015. Washington DC: World Bank (2018) http://sdwebx.worldbank.org

Young G, Angier P. Developing a fair trade certification label for rice exports from Guyana & Suriname. Capetown: Imani Development (2010). 55 p.

Zeven AC. Landraces: a review of definitions and classifications. Euphytica (1998) 104:127–39. https://doi.org/10.1023/A:1018683119237

