## Supplementary Material for "Hidden Rice Diversity in the Guianas"

**Links to video footage made on Maroon rice cultivation and processing.**

Video 1. Two ways of planting rice: either in small holes (‘diki olo’) dug with a hoe (‘tyap’), or cast over a cleaned field (‘fringi’) after which the rice is covered with loose soil and pressed flat.

<https://www.youtube.com/watch?v=rWUDuZCKFr0>

Video 2. Harvesting rice with a small knife in St. Laurent du Maroni, French Guiana. This video also shows how interviews were carried out in the field.

<https://www.youtube.com/watch?v=bQ1tyfhXIJI&t=110s>

Video 3. Traditional way of threshing rice by Maroons in French Guiana. <https://www.youtube.com/watch?v=srwL5GrLLW4>

Video 4. Traditional dehusking of rice by Maroons in Suriname.

<https://youtu.be/KWXIXHhhf4g>

Video 5. Traditional winnowing of rice with a wooden tray in Suriname.

<https://www.youtube.com/watch?v=_5dbhOnqAMU&feature=youtu.be>

Video 6. How the Maroon ancestors hid rice grains in their hair.

<https://www.youtube.com/watch?v=4H1IbY6PGIk>

**Supplementary Figures**

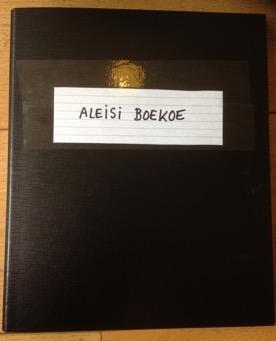
  
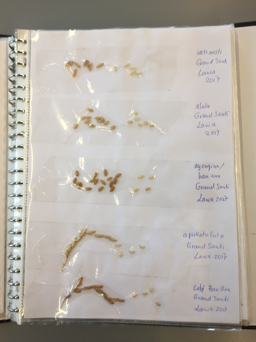

**Supplementary Figure S1.** A. Ring binder named ‘aleisi boekoe’ (rice book in Sranantongo). Picture by Tinde van Andel. B. Pages containing different rice varieties. Picture by Tinde van Andel.

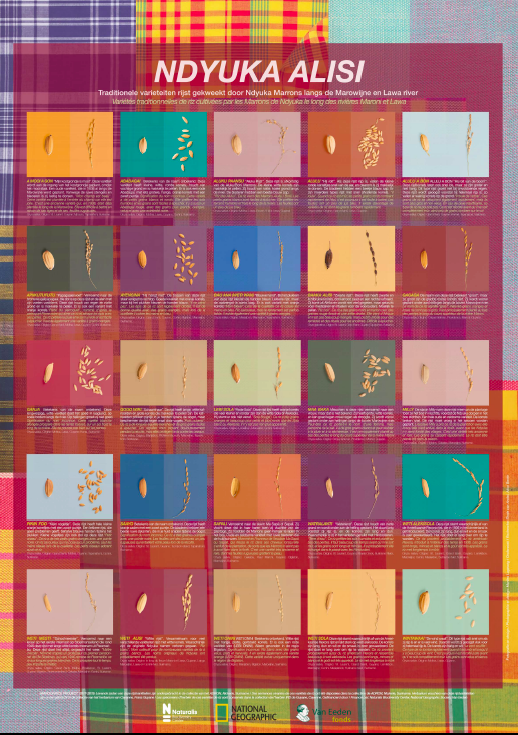

**Supplementary Figure S2**. Poster with pictures of the different rice varieties and associated traditional information in Dutch and French to be distributed in Suriname and French Guiana. Photography and design by Marlies Lageweg.

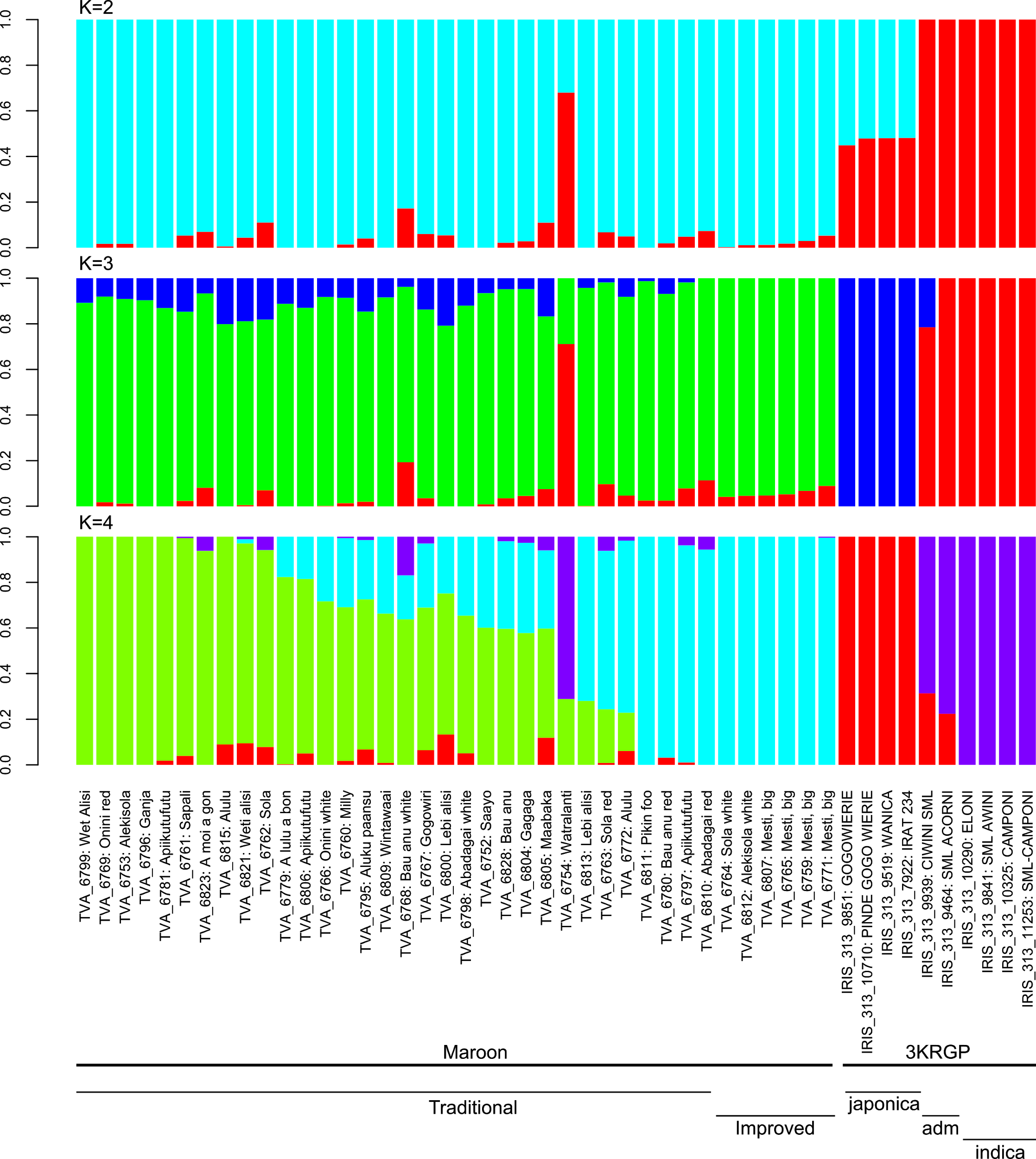

**Supplementary Figure S3.** Population structure results for all primary (Maroon) and secondary (3KRGP) *O. sativa* accessions, based on 64,313 SNPs. A. Cross-validation error estimates demonstrate that the optimal number of ancestral populations given the data is K=2.

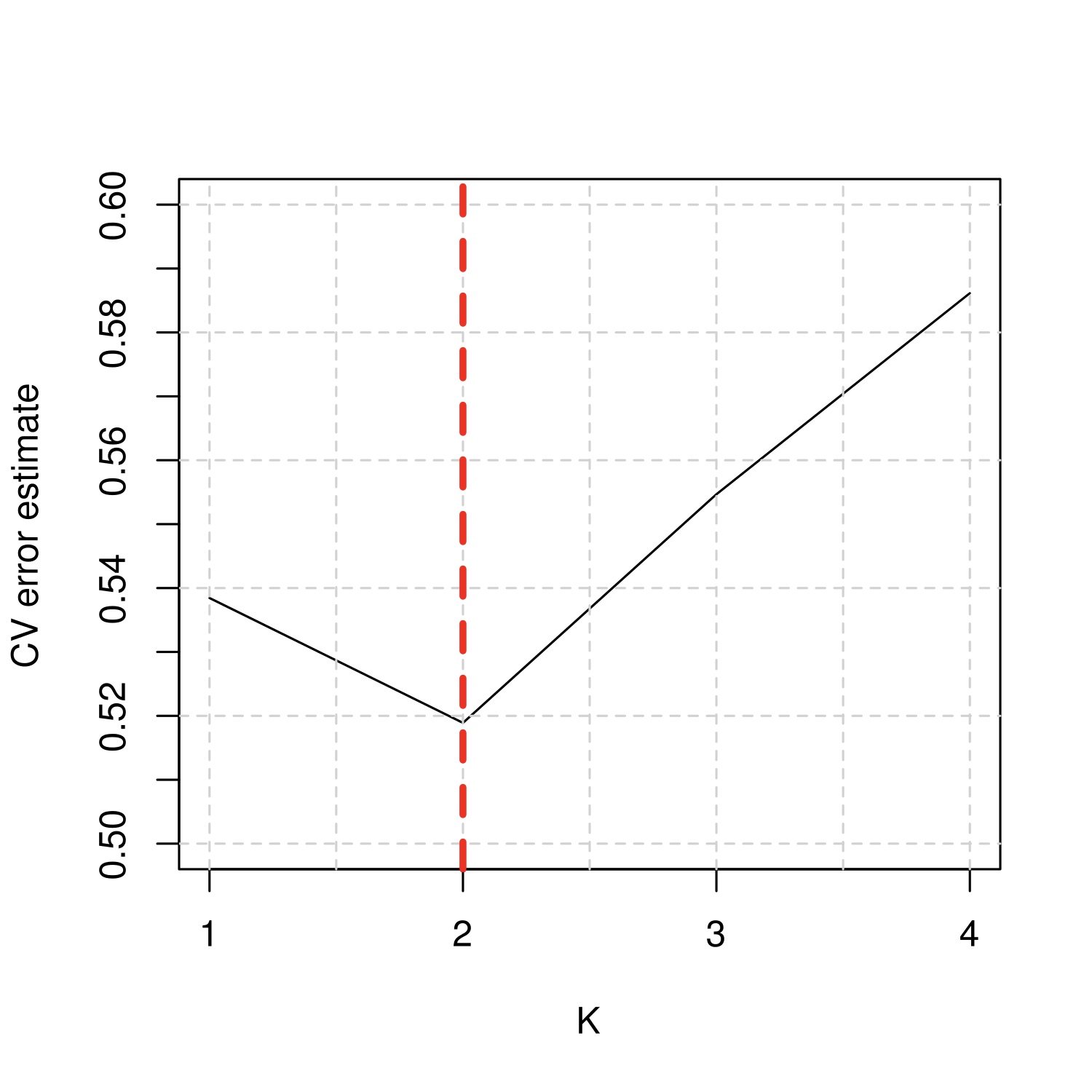

**Supplementary Figure S3.** Population structure results for all primary (Maroon) and secondary (3KRGP) *O. sativa* accessions, based on 64,313 SNPs. **B.** Relative ancestry fractions per individual at different levels of K.

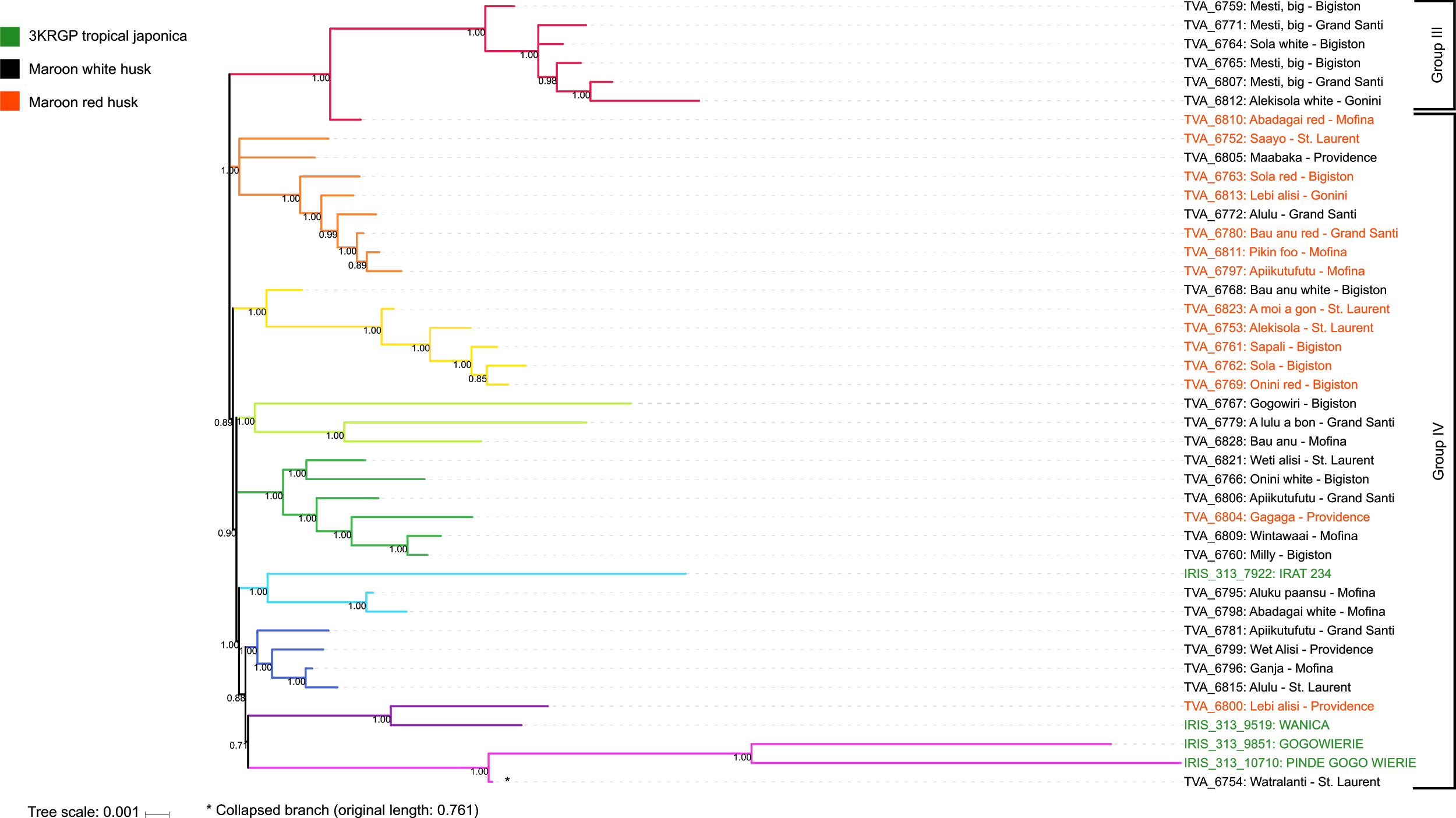

**Supplementary Figure S4.** Clustering of the Maroon Asian rice gene pool. A. Approximately ML tree based on 39,595 homozygous SNPs, with branch lengths representing evolutionary distances between all Maroon and 3KRGP tropical japonica varieties. The branch connecting to the sole wetland variety is collapsed for visualization purposes and marked with an asterisk. Clades with good support values (>0.85) on all internal nodes have been identified and labelled with different colors.

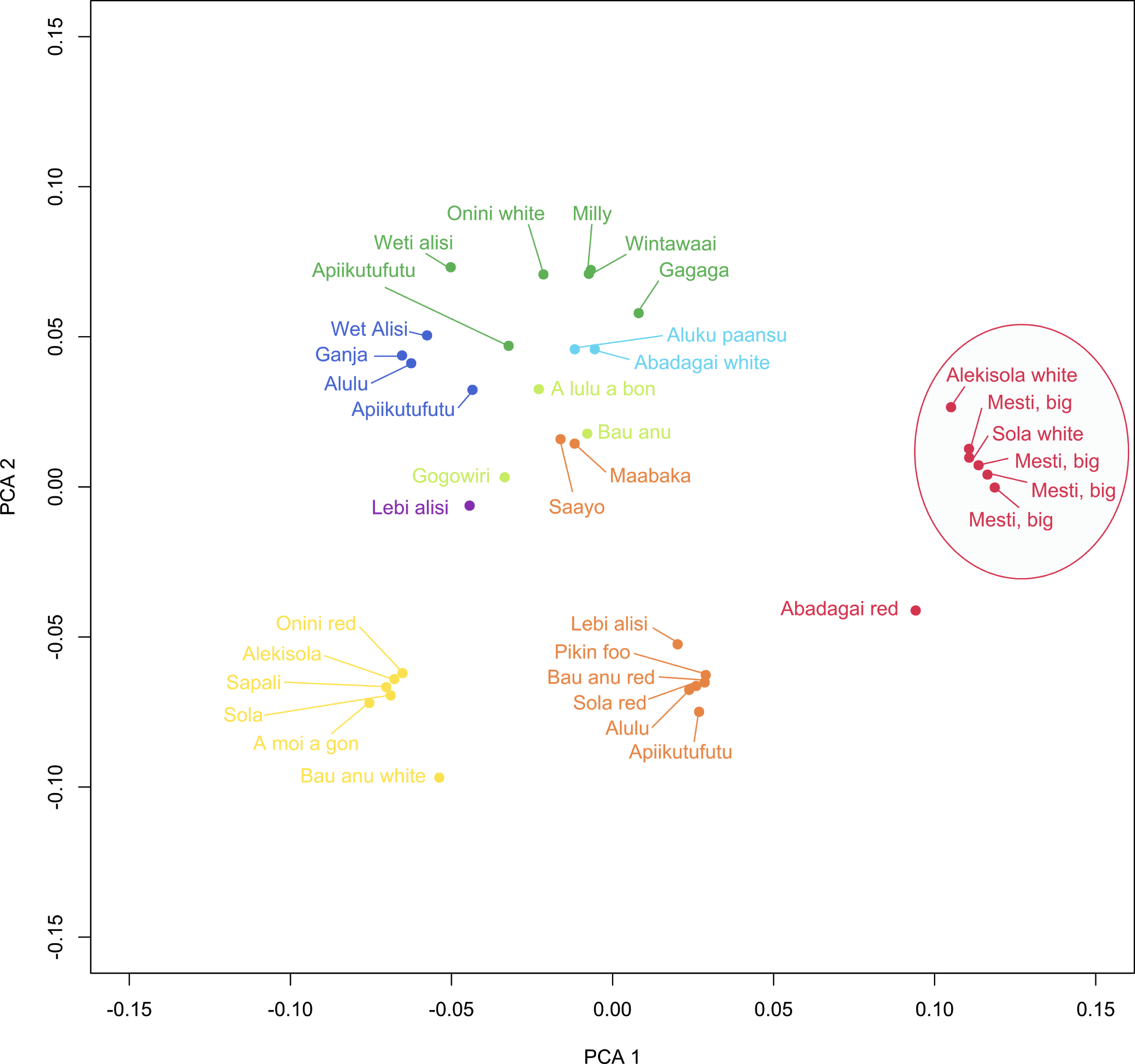

**Supplementary Figure S4.** Clustering of the Maroon Asian rice gene pool. B. Multi-dimensional scaling in two dimensions of raw Hamming distances for all improved (Group III) and traditional dryland (Group IV) varieties. Individuals are labelled by vernacular name and color-coded based on the clustering in **Figure S4A.** Improved varieties are circled.
